# Highly enriched hiPSC-derived midbrain dopaminergic neurons robustly models Parkinson’s disease

**DOI:** 10.1101/2020.09.08.287797

**Authors:** Gurvir S Virdi, Minee L Choi, Zhi Yao, James R Evans, Dilan Athauda, Daniela Melandri, Sergiy Sylantyev, Andrey Y Abramov, Rickie Patani, Sonia Gandhi

## Abstract

The development of human induced pluripotent stem cells (hiPSC) has greatly aided our ability to model neurodegenerative diseases. However, generation of midbrain dopaminergic (mDA) neurons is a major challenge and protocols are variable. Here, we developed a method to differentiate hiPSCs into enriched populations (>80%) of mDA neurons using only small molecules. We confirmed the identity of the mDA neurons using single-cell RNA-sequencing and detection of classical markers. Single-cell live imaging demonstrated neuronal calcium signalling and functional dopamine transport. Electrophysiology measures highlighted the ability to form synapses and networks in culture. Patient-specific hiPSC lines differentiated to produce functional mDA neurons that exhibit the hallmarks of synucleinopathy including: aggregate formation, oxidative stress as well as mitochondrial dysfunction and impaired lysosomal dynamics. In summary, we establish a robust differentiation paradigm to generate enriched mDA neurons from hiPSCs, which can be used to faithfully model key aspects of Parkinson’s disease (PD), providing the potential to further elucidate molecular mechanisms contributing to disease development.

## Introduction

Parkinson’s disease (PD) is a progressive neurodegenerative disease characterised primarily by the loss of dopaminergic neurons in the substantia nigra in the midbrain (Poewe *et al*., 2017). Studying the mechanisms that contribute to disease is of immense importance, but has been somewhat limited by a paucity of relevant human models. The development of human induced pluripotent stem cells (hiPSCs) from patients with familial forms of PD has vastly improved our capability to model and study PD and offers a complimentary approach in a more relevant human model. Following developmental cues, hiPSCs can be patterned into region-specific parts of the brain, including the midbrain.

During *in vivo* development, gradients of various morphogens result in the activation of different key transcription factors, which dictate, and define the cytoarchitecture of different regions of the brain. Upon the formation of the neural tube, midbrain dopaminergic (mDA) neurons, which form the substantia nigra emerge from the ventral mesencephalic floor plate (Marín *et al*., 2005; Ono *et al*., 2007). Their development is dependent on key signalling centres including the floor plate and the isthmic organiser. At the floor plate, a high concentration of the morphogen Sonic hedgehog (SHH) is established, which is essential for mDA neurogenesis (Arenas *et al*., 2015). In addition, mDA neurogenesis also depends on the presence of the key morphogens, Wnt1 (wingless-int1), and FGF8 (fibroblast growth factor 8) from the isthmic organiser, located at the midbrain/hindbrain boundary. Together, these morphogens provide the essential cues to induce developmental programmes that ‘pattern’ the cells to the ventral mesencephalon (mDA progenitors), which subsequently differentiate to yield mDA neurons (Arenas *et al*., 2015).

Since the discovery of hiPSCs, several protocols have been developed to specify midbrain dopaminergic (mDA) neurons, the cell type primarily vulnerable in PD (Doi *et al*., 2014; Kirkeby *et al*., 2012; Kriks *et al*., 2011; Nolbrant *et al*., 2017). These protocols mimic *in vivo* developmental programmes, inducing neuronal induction via dual-SMAD inhibition (Chambers *et al*., 2009), and activating SHH, Wnt, and FGF signalling through the use of small-molecules and/or recombinant morphogens. The resulting cells are molecularly defined based on the expression of key genes and proteins, as well as functional and physiological characteristics resembling *in vivo* mDA neurons (Hartfield *et al*., 2014). However, the efficiency of mDA neuron production is highly variable (10-70%) following current differentiation protocols (Marton and Ioannidis, 2019). Furthermore, differentiation protocols are often expensive due to the use of recombinant protein factors, to pattern and differentiate cells, and take a long time to yield high purity mDA neurons.

At a molecular level, the insoluble aggregated forms of alpha-synuclein (coded by the *SNCA* gene) are a common hallmark of sporadic, and familial forms of PD (Choi and Gandhi, 2018). Using hiPSC-derived mDA neurons to model alpha-synuclein pathology is thus vital to increase our understanding of the underlying mechanisms of pathology. Generation of a hiPSC line from a patient with early onset PD, and a triplication in the *SNCA* locus was shown to have elevated levels of the alpha-synuclein protein (Devine, Ryten, *et al*., 2011). We have previously shown that the triplication of *SNCA* locus results in an increase in alpha-synuclein oligomers, which interact with ATP synthase, causing mitochondrial dysfunction and resulting in the opening of the mitochondrial transition permeability pore (PTP), which causes neuronal death (Ludtmann *et al*., 2018). Similarly, a point mutation in SNCA (A53T) causes early onset, progressive PD (Spira *et al*., 2001). A53T hiPSC-derived neurons have been shown to have an increase in nitrostative, and ER stress (Chung *et al*., 2013), as well as an increase in endogenous alpha-synuclein and an increase in mitochondrial dysfunction (Zambon *et al*., 2019).

As current protocols result in terminally differentiated mDA cultures of variable enrichment (Marton and Ioannidis, 2019), this often hampers the ability to identify cell type specific phenotypes. Recently, enrichment of only tyrosine hydroxylase (TH) positive hiPSC-derived mDA neurons has discovered new disease mechanisms in glucocerebrosidase (GBA) linked PD (Lang *et al*., 2019), highlighting the need for higher enrichment of mDA cultures. Thus, we aimed to establish a protocol to obtain highly enriched mDA neurons and test their capacity to faithfully model key aspects of PD pathogenesis as well as identify early phenotypes. We developed a protocol to differentiate hiPSCs into highly enriched mDA neurons (>80%), which are functional and electrically mature. We demonstrate this is a relevant model for PD pathology, as A53T and *SNCA* triplication lines displayed common alpha-synuclein related pathology. Moreover, our protocol is more economically favourable as it does not rely on recombinant proteins to differentiate hiPSCs to mDA neurons.

## Materials and Methods

Detailed material and methods can be found in the supplementary material

### Human induced pluripotent stem cell culture

Human induced pluripotent stem cells (hiPSCs) were maintained in feeder-free monolayers on Geltrex (ThermoFisherScientific) and fed daily with Essential 8 medium (Life Technologies). When confluent, hiPSCs were passaged using 0.5 μM EDTA (Life Technologies). All cells were maintained at 37°C and 5% carbon dioxide.

### Data availability

The data that supports the findings of this study is available by contacting the corresponding authors upon reasonable requests.

## Results

### Generating an enriched population of mDA neurons from hiPSCs

To generate mDA neurons, we first treated hiPSC lines from healthy controls with the previously reported dual SMAD inhibition (Chambers *et al*., 2009) for neural conversion. Lineage restriction towards the midbrain was achieved by treating our cultures with small molecule agonists of SHH and WNT, purmorphamine and CHIR99021 respectively (Fig. 1A). After 14 days of the induction, we confirmed mDA neural precursor cell (NPC) identity by first, profiling the expression of mDA NPC marker genes LMX1A, FOXA2, and EN1 by RT-PCR, which revealed clear increases in expression (Fig. 1B) (p < 0.005). We then demonstrated an increased protein expression of the midbrain NPC markers LMX1A and FOXA2, as well as the diencephalon marker OTX2 using immunocytochemistry (ICC) (Fig. 1C). Quantification of cells co-expressing all three markers showed an enriched culture of mDA NPCs of approximately 85% across all lines (Fig. 1D) (Ctrl 1: 83% ± 3.5, Ctrl 2: 81.4% ± 2.4, Ctrl 3: 88.7% ± 3.1, Ctrl 4: 88.4% ± 2.2). After midbrain patterning, NPCs were maintained in culture for four days before terminal differentiation, in order to increase their confluence. Differentiation of NPCs was achieved using a Notch pathway inhibitor and a Rho-associated protein kinase (ROCK) inhibitor, encouraging cell cycle exit and cell survival respectively. The expression of the mature mDA marker TH increased during terminal differentiation, achieving approximately 75% of TH positive cells after 3 weeks (p < 0.0001) (Fig. 1E). At this stage, we quantified the proportion of TH positive cells using two approaches. Firstly, quantification via ICC showed that approximately 75-80% of DAPI positive cells expressed TH (Fig. 1F, 1G) (Ctrl 1: 82.1% ± 3.8, Ctrl 2: 73.1% ± 6.9, Ctrl 3: 77.9% ± 3.2). Secondly, using flow cytometry (see materials and methods section), we found that the proportion of cells expressing both TH and β-III Tubulin was measured to be approximately 88% (Fig. 1H, Fig. 1I) (Supplementary Fig 1A, 1B) (88.8% ± 1.9). Overall the data confirms high differentiation efficiency into neurons, and specifically into TH positive neurons, up to 80-90% of the whole culture, with comparable efficiencies across 3 control hiPSC lines. To verify the maturity of the TH positive mDA neurons, qPCR for mDA specific genes (nuclear receptor related-1 protein (Nurr1), G protein-activated inward rectifier potassium channel 2 (GIRK2), and the dopamine transporter (DAT)) was performed in mDA neurons 3-4 weeks post terminal differentiation. We demonstrated an increase in the expression of all these genes relative to midbrain NPCs (Supplementary Fig. 1C). GIRK2 expression was further confirmed by ICC using a knockout validated antibody. GIRK2 is highly expressed in dopaminergic neurons in the A9 region of the midbrain (typical of the substantia nigra), confirming the identity of the TH positive neurons further (Supplementary Fig. 1D). Interestingly, we observed a nuclear and cytosolic signal using the GIRK2 antibody, both of which were abolished on applying the antibody incubated with the control GIRK2 antigen (Supplementary Fig. 1E).

**Figure 1.**
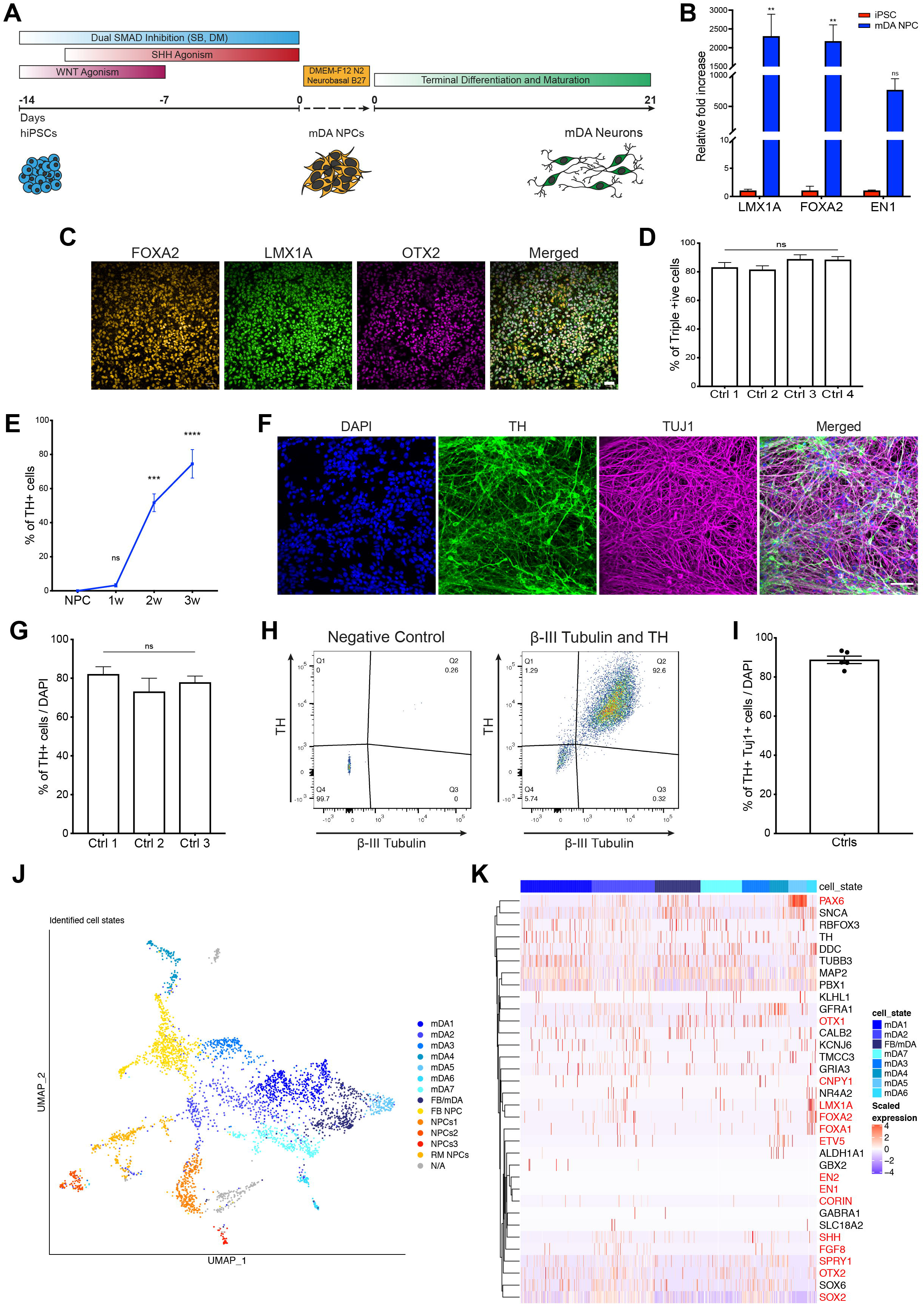
Characterisation of an enriched population of mDA NPCs and neurons. **(A)** Illustration depicting the differentiation protocol to generate mDA neurons. The hiPSCs are patterned to midbrain NPCs after 14 days. NPCs are then maintained in culture for 4 days before being terminally differentiated to yield mature mDA neurons after 21 days. **(B)** Quantitative PCR showing a high up-regulation of mRNA for mDA NPC markers LMX1A, FOXA2, and EN1 relative to hiPSCs (n= 3 different lines across 4 independent neuronal inductions, ns p > 0.05 ** p < 0.005, ordinary two-way ANOVA). Data plotted as ±SEM. **(C)** Representative immunocytochemistry images showing high expression of mDA NPC markers FOXA2, LMX1A, and the forebrain/midbrain marker OTX2 (scale bar = 50μm). **(D)** Quantification of immunocytochemistry images showing >80% of cells co-express mDA NPC markers FOXA2, LMX1A, and OTX2 (n = 4 different lines across 3 independent neuronal inductions, ns p > 0.05, ordinary one-way ANOVA). Values plotted as ±SEM. **(E)** Quantification of immunocytochemistry images showing the increase in expression of the mDA mature neuron marker, TH, as the neurons differentiate and mature (n = 3 different lines across 1 independent neuronal induction, ns p > 0.05, *** p < 0.0005, **** p < 0.0001, ordinary one-way ANOVA). Values plotted as ±SEM. **(F)** Representative immunocytochemistry images showing the expression of TH and the neuronal marker, TUJ1 after 3 weeks of differentiation (41 days from hiPSC) (scale bar = 50μm). **(G)** Quantification of immunocytochemistry images after 3 weeks of differentiation showing approximately 80% of total cells express TH (n= 3 different lines across 3 independent neuronal inductions, ns p > 0.05, ordinary one-way ANOVA). Values plotted as ±SEM. **(H)** Representative dot plots of single-cell suspensions showing the percentage of TH and β-III Tubulin positive cells in the mDA cultures 3 weeks after terminal differentiation. A negative control (DAPI only) was used to determine quantification thresholds to set the gating (n = 10,000 events recorded per measurement). **(I)** Quantification of flow cytometry analysis showing over 80% of DAPI positive cells co-express TH and β-III Tubulin (n = 3 lines across 2 independent neuronal inductions). Values plotted as ±SEM. **(J)** A UMAP plot showing the 13 clusters identified from single-cell RNA-seq after 4 weeks of differentiation. Neuronal mDA (mDA1-7), and forebrain/midbrain neuron (FB/mDA) clusters are coloured in blue, and mDA NPC/neurons (NPCs1-3), forebrain NPCs (FB NPCs), and rostral midbrain NPCs (RM NPCs) are coloured in red, orange, and yellow. An unidentified cluster (N/A) is coloured in grey. **(K)** A heat map showing the expression of genes in clusters identified as mDA neurons (mDA1-7), and the forebrain/midbrain neuron (FB/mDA) cluster in the culture after 4 weeks of differentiation. Each line represents a cell from that cluster. Genes annotated in red correspond to mDA NPC markers, and those in black correspond to mDA neuron markers.

In order to further validate neuron identity, we performed single-cell RNA-seq on mDA neurons after 4 weeks of differentiation (day 48 from hiPSC). The cells were clustered into identities based on their transcriptional profiles. Clustering of all grouped control mDA neurons revealed 13 clusters, of which 7 expressed key mDA neuron genes, suggesting they are mDA neuron in identity (Fig. 1J) (Supplementary Fig. 2A). The 7 mDA neuron clusters expressed key mDA genes including GDNF receptor (GFRA1), NURR1, SOX6, PBX1, TH, GIRK2, NURR1, DDC, which suggests that most of the cells captured are of mDA identity, and are abundant within the culture (Fig. 1K). Interestingly, within the data we also detected several clusters which appeared to be similar to mDA NPCs, or early neurons (5 clusters) (Supplementary Fig. 2B). These clusters expressed key NPC markers like FOXA2, LMX1A, OTX2, FGF8, SHH, and also expressed mDA neuron markers like GFRA1, DDC, NURR1, TH, GIRK2 (Supplementary Fig. 2B, 2C). In addition, most clusters expressed *SNCA* highlighting the potential to use the protocol as a model of synucleinopathy (Supplementary Fig. 2D). Taken together, our analysis suggests that the majority of the cell culture are mDA neuron in identity, or mDA NPC like, on course to differentiate to mDA neurons.

**Figure 2.**
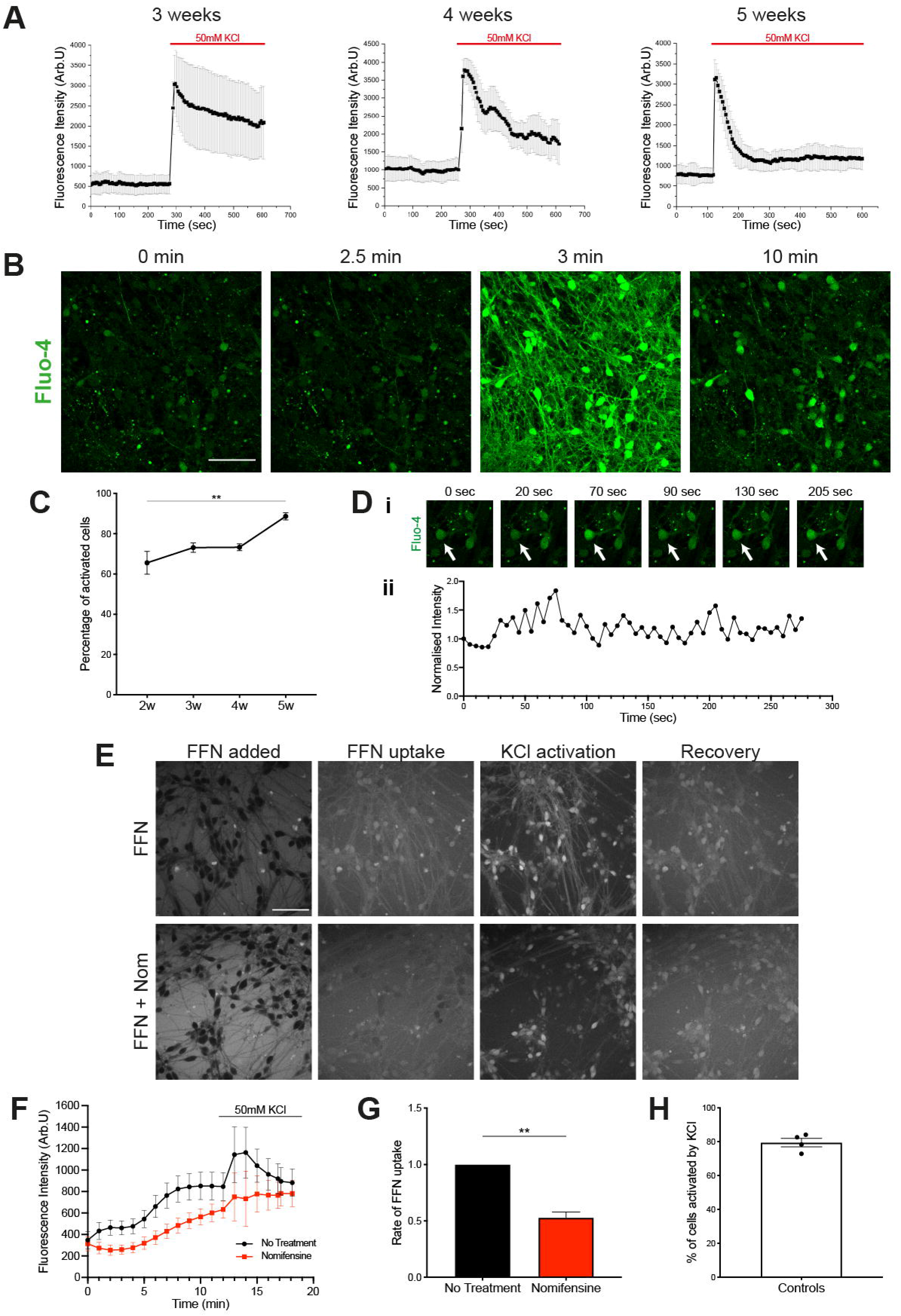
Functional characterisation of mDA neurons. Representative traces of the time series showing the fluorescence intensity of Fluo-4 increase and recover upon 50mM KCl addition at 3, 4, and 5 weeks of terminal differentiation (n = 15 cells per line, 3 separate lines across 1 independent neuronal induction). Values plotted as ±SD. **(B)** Representative pictures of mDA neurons at 5 weeks of terminal differentiation during a time series, showing the levels of intracellular calcium using the fluorescent calcium indicator dye Fluo-4 (scale bar = 50μm). **(C)** Quantification of the number of cells that had a calcium spike upon 50 mM KCl stimulation across different weeks of terminal differentiation (n = 3 separate lines across 1 independent neuronal induction, ns p > 0.05, ** p = 0.0075, two-way ANOVA). Values plotted as ±SEM. **(D)** Example of spontaneous calcium activity at 3 weeks of terminal differentiation showing, **(Di)** pictures at different times prior to KCl stimulation, arrow indicates the highlighted cell from which, **(Dii)** recordings of the fluorescence change in Fluo-4 intensity was recorded. **(E)** Representative images showing the uptake of the DAT fluorescent substrate FFN102 (FFN) in mDA neurons 4-5 weeks of terminal differentiation. The first panel shows addition of FFN dye where it is outside of the cells. Second panel shows the dye inside the cells. Third panel shows the response of the dye upon stimulation with 50mM KCl and the recovery in the fourth panel. Lower panels show the dye behaviour in the presence of the DAT inhibitor, nomifensine (scale bar = 50μm). **(F)** Representative trace showing the increase in intracellular fluorescence of FFN as time increases, as well as response to KCl stimulation, highlighting DAT presence and function. The DAT inhibitor nomifensine reduces the rate of FFN dye uptake showing DAT specificity (n = 15-20 cells per condition). Values plotted as ±SD. **(G)** Quantification of the normalised rate of FFN uptake and the rate of uptake in the presence of nomifensine (n = 3 separate lines over 3 independent neuronal inductions, Welch’s t-test, ** p = 0.0024). Values plotted as ±SEM. **(H)** Quantification of the number of FFN positive cells once stimulated with the addition of KCl (n = 3 separate lines, 3 independent neuronal inductions). Values plotted as ±SEM.

### Functional characterisation of differentiated mDA neurons

In order to test the functional properties of the neurons, we first determined whether they expressed voltage gated calcium channels, which are expressed in neurons and not astrocytes. Opening of calcium channels is achieved using a high concentration of KCl (50mM), which depolarises the plasma membrane resulting in a large influx of calcium into the neurons. Neurons were loaded with the calcium indicator Fluo4-AM, and 50mM KCl was applied to the culture (Fig. 2A, Fig. 2B) (Supplementary Fig. 3A). We demonstrated a Ca^2+^ signal induced by KCl in all cultures as early as 2 weeks post differentiation. The proportion of cells responding to KCl was 65% after 2 weeks, increasing to over 85% after 5 weeks of differentiation (Fig. 2C) (2 weeks = 65.5% ± 5.7, 3 weeks = 73.1% ± 2.3, 4 weeks = 73.2 ± 1.7, 5 weeks = 88.6% ± 1.8, p = 0.0075). Furthermore, we observed the calcium efflux, as shown by the decay of the calcium signal, was significantly altered over time in culture, presumably reflecting maturation of the efflux pumps (Fig. 2A) (Supplementary Fig. 3B). A major characteristic of mature neurons is their spontaneous calcium activity. Oscillations of Ca^2+^ have been shown to be a key hallmark of mDA dopaminergic neurons in the substantia nigra, which contributes to their pace-making characteristic (Guzman *et al*., 2009). We similarly observed spontaneous calcium fluctuations in neurons after 3 weeks of differentiation, as shown using single cell imaging of fluo-4 labelled neurons (Fig. 2D). This suggests that our mDA neurons are functional, and spontaneously oscillate calcium.

**Figure 3.**
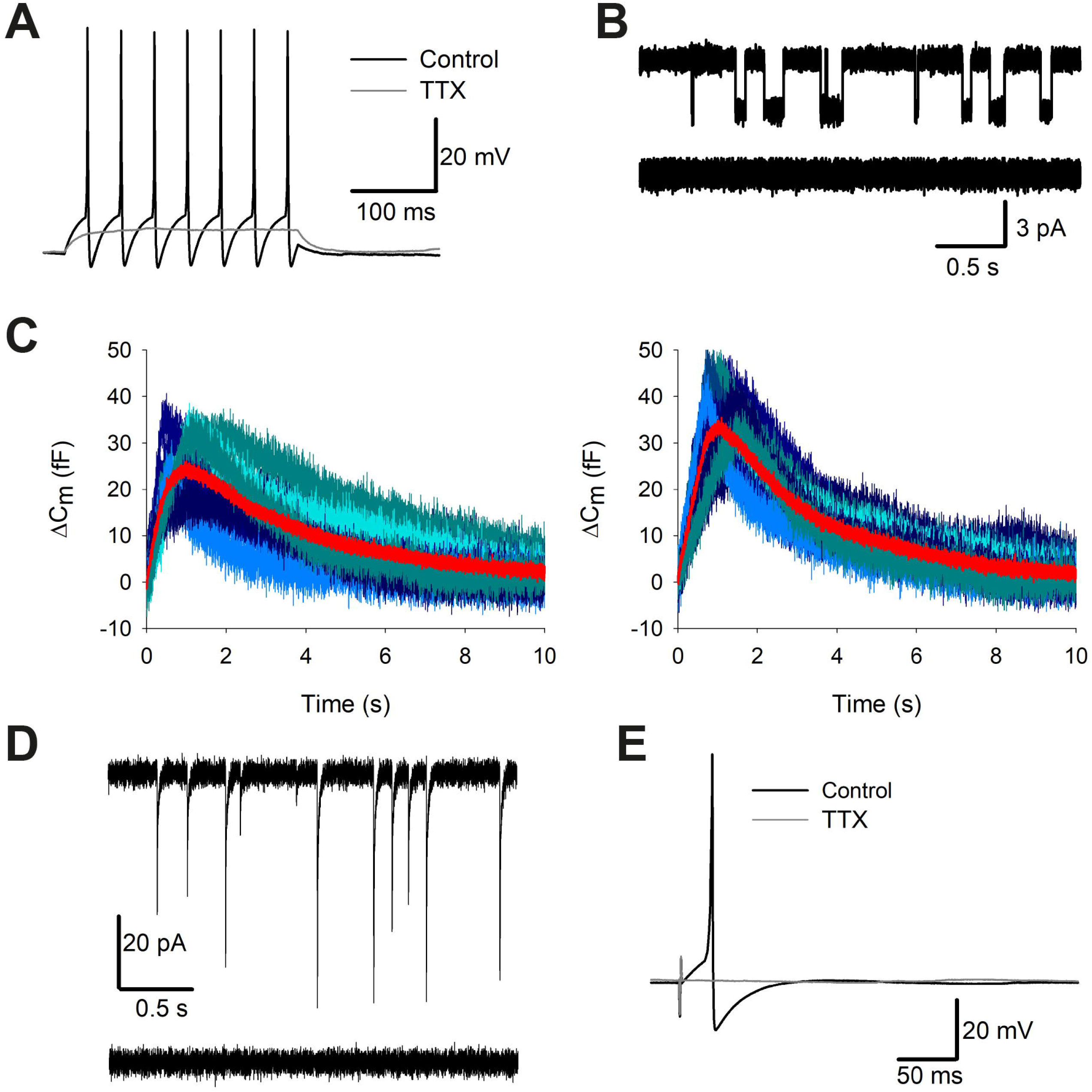
Electrophysiological characterisation of mDA neurons. **(A)** APs triggered by step current injection in mDA neurons at day 30 of differentiation. 1 mM tetrodotoxin (TTX) fully suppresses APs, confirming involvement of voltage-gated sodium channels. **(B)** Single-channel openings of NMDA receptors in an outside-out patch excised from mDA neurons at day 30 of differentiation. Top trace: application of 10 mM glutamate + 10 mM glycine triggers single-channel openings with two different conductance states. Bottom trace: 50 mM APV fully suppresses effect of glutamate + glycine, confirming pharmacological profile of NMDA receptors. **(C)** Changes in whole-cell membrane capacitance evoked by stimulation series confirm elevated intensity of vesicle release in mDA neurons at day 70 of differentiation. Shadows of blue, high noise: 10 consequent individual traces; superficial red trace with low noise: averaged trace. Left: control. Right: elevated Ca^2+^ magnifies the effect of external stimulation on membrane capacitance, thus confirming involvement of presynaptic Ca^2+^-dependent mechanism of vesicle release. **(D)** Spontaneous postsynaptic activity confirms presence of functional synapses in mDA neurons at day 70 of differentiation. Top trace: spontaneous postsynaptic currents. Bottom trace: 50 mM of picrotoxin fully suppresses spontaneous synaptic activity, confirming its generation by GABA_A_ receptors. **(E)** AP generation in response to field stimulation confirms establishment of a functional neuronal network in mDA neurons at day 105 of differentiation.

We next investigated the function of the DAT by measuring the uptake of the fluorogenic DAT substrate FFN102 (FFN), which enters cells through DAT (Rodriguez *et al*., 2013). When incubated with the dye, followed by a wash, the dye can be seen inside mDA neurons (Supplementary Fig. 3C). We show a basal uptake of FFN in neurons which is further activated upon KCl induced calcium influx in the majority of neurons (Fig. 2E, Fig. 2H) (cells activated by KCl = 79.5% ± 2.5). Inhibition of the DAT using the compound nomifensine resulted in a significantly reduced basal uptake rate of FFN into neurons (No treatment = 1.00 ± 0.00, Nomifensine = 0.53 ± 0.05, p = 0.0024), as well as a reduced KCl stimulated spike (Fig. 2F, Fig. 2G) (Supplementary Fig. 3D). This suggests that our mDA neurons express functional DAT.

### Electrophysiological characterisation of differentiated mDA neurons

To obtain functional electrophysiological characteristics of the generated mDA neurons, we performed a series of experiments, targeting single-channel, single-cell and neuronal network effects (Fig. 3). Our first experiment confirmed the presence of action potential (AP) generation machinery activated by depolarizing voltage step, with the classical tetrodotoxin (TTX)-sensitive voltage-gated sodium channel involvement (Fig. 3A). Secondly, we confirmed the presence of excitatory NMDA receptors. 10mM glutamate + 10mM glycine added to the perfusion solution triggered single-channel openings in outside-out patches pulled from cell somata; further application of 50mM APV (selective antagonist of NMDA receptors) fully suppressed this effect (Fig. 3B). Thirdly, we tested whether our cultured neurons have a functional presynaptic neurotransmitter release system. To do this, we recorded changes in cell membrane capacitance (C_m_) in response to a field stimulation train of five stimuli with 50ms intervals (Fig. 3C). To test the involvement of the classical Ca^2+^-dependent mechanism of presynaptic vesicle release, we repeated the experiment under normal (2 mM) and elevated (5 mM) Ca^2+^ concentration. Here we observed a clear increase between C_m_ value under normal Ca^2+^ concentration and after an elevated Ca^2+^ concentration (Fig. 3C), confirming Ca^2+^-dependent presynaptic vesicle release. Next, we performed a whole-cell voltage-clamp recording of spontaneous postsynaptic activity. In this experiment we observed postsynaptic currents, which were suppressed by 50mM of the selective antagonist of GABA_A_ receptor, picrotoxin (Fig. 3D). This confirmed the presence of the main type of inhibitory neuroreceptors (GABA_A_ receptors) and of a functional postsynaptic system with a classical response to spontaneously released GABA. Finally, we set out to test whether our mDA neurons can establish a functional neural network. To achieve this, we performed whole-cell current-clamp recordings, delivering field electrical stimuli in a perfusion chamber. These stimuli generated APs of classical shape, confirming the presence of interneuronal crosstalk (Fig. 3E). Thus, we show that the mDA neurons, by week 15 of differentiation, display many electrophysiological characteristics that confirm their behaviour as neurons forming functional networks (summarised in Table 1).

**Table 1.** Electrophysiological effects observed in mDA neurons.

| Electrophysiological effect | Ctrl 1 | Ctrl 2 | Ctrl 3 | SNCA x3 |
| --- | --- | --- | --- | --- |
| AP generation in response to voltage step | X | X | X | X |
| Single-channel NMDA receptor activity | X | X |  | X |
| Stimulation-evoked changes of $C_m$ | | X | X | X |
| GABA-ergic spontaneous activity |  |  | X |  |
| APs evoked by field stimulation | X | X |  |  |

### Patient-specific hiPSCs differentiate into functional mDA neurons

Mutations in the gene encoding alpha-synuclein, *SNCA*, cause an autosomal dominant form of PD (SNCA PD). We obtained hiPSCs from two patients with the A53T mutation, and a patient with a triplication of the *SNCA* locus (SNCA x3), and differentiated them into midbrain dopaminergic neurons according to the protocol described above. Both of these genetic changes cause an early onset and rapidly progressive form of PD (Devine, Gwinn, *et al*., 2011); (Farrer *et al*., 2004); (Spira *et al*., 2001). All three mutant lines were successfully differentiated into mDA neurons, highlighted by a high expression of TH and MAP2 (Fig. 4A). We next functionally validated SNCA neurons by confirming that at 2 weeks post differentiation, 60-70% of neurons exhibit a cytosolic calcium response to KCl confirming the presence of voltage gated calcium channels (Fig. 4B, Fig. 4C) (Control = 65.5% ± 5.7, A53T = 71.9% ± 1.7, SNCA x3 = 66%). SNCA neurons exhibited TTX-sensitive action potential generation in response to injected current (Fig. 4D), single channel NMDA receptor activity (Figure 4E), and stimulation evoked changes in capacitance (Fig. 4F), confirming characteristic electrophysiological activity in patient-specific mDA neurons.

**Figure 4.**
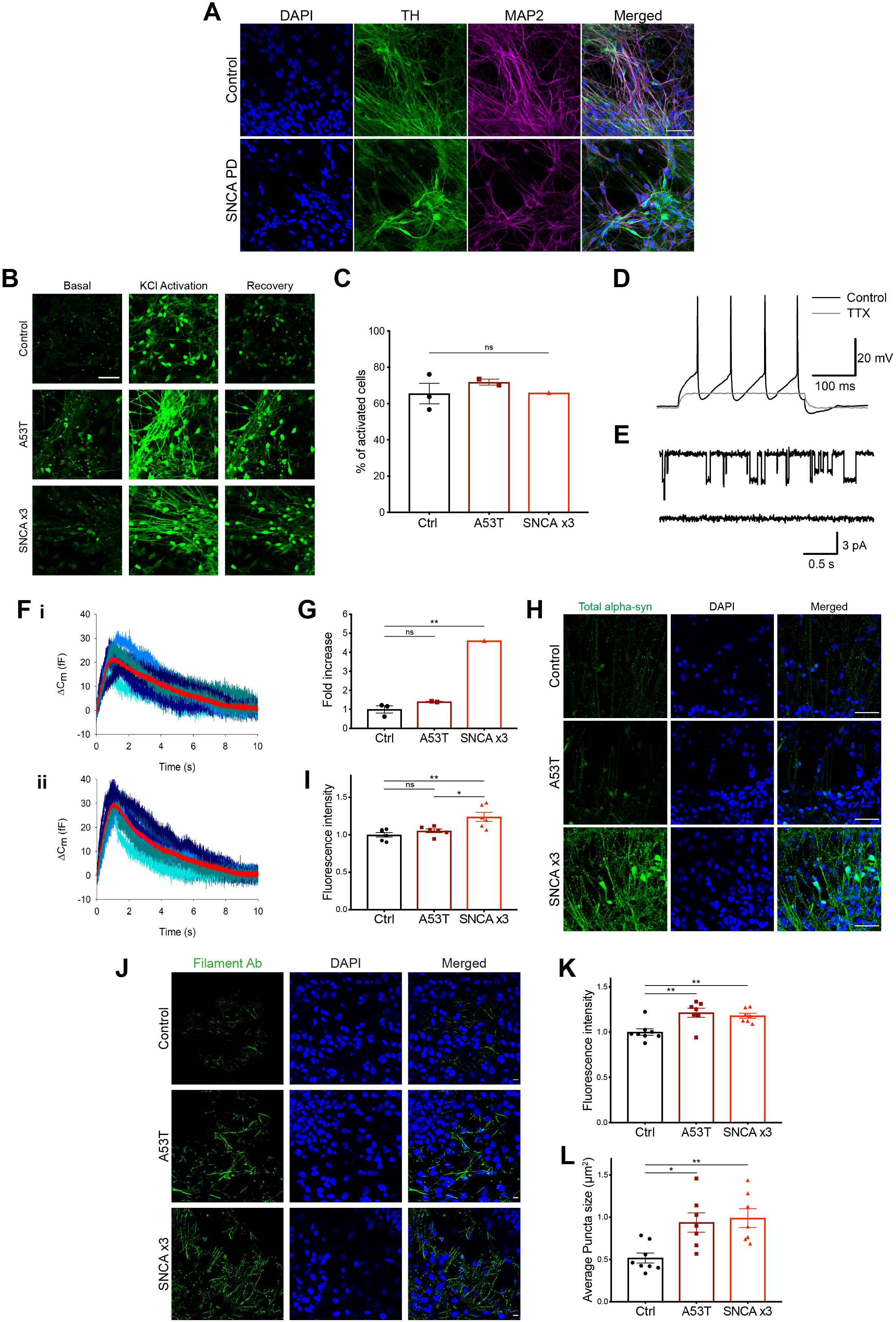
Generation and characterisation of mDA neurons from hiPSC lines from patients with alpha-synuclein mutations. **(A)** Representative immunocytochemistry images of control and alpha-synuclein (SNCA) patient hiPSCs showing the expression of TH and MAP2. Scale bar = 50μm. **(B)** Representative time series snaps of Fluo-4 Ca^2+^ recordings in control, A53T, and SNCA x3 lines showing basal intensity as well as the response to 50mM KCl stimulation, and the recovery at 2 weeks of terminal differentiation (scale bar = 50μm). **(C)** Quantification of the number of cells that had a calcium spike upon KCl stimulation after 2 weeks of differentiation (n = 3 control hiPSC lines, 2 A53T patient lines, 1 SNCA x3 patient line, 1 neuronal induction, ns p > 0.05, one-way ANOVA). Values plotted as ±SEM. **(D)** APs triggered by step current injection in SNCA x3 mDA neurons at day 70 of differentiation. 1 mM tetrodotoxin (TTX) fully suppresses APs, confirming involvement of voltage-gated sodium channels. **(E)** Single-channel openings of NMDA receptors in an outside-out patch excised in SNCA x3 mDA neurons at day 70 of differentiation. Top trace: application of 10 mM glutamate + 10 mM glycine triggers single-channel openings with two different conductance states. Bottom trace: 50 mM APV fully suppresses effect of glutamate + glycine, confirming pharmacological profile of NMDA receptors. **(F)** Changes in whole-cell membrane capacitance evoked by stimulation series confirm elevated intensity of vesicle release in SNCA x3 mDA neurons at day 70 of differentiation. Shadows of blue, high noise: 10 consequent individual traces; superficial red trace with low noise: averaged trace. i) control conditions. ii) elevated Ca^2+^ magnifies the effect of external stimulation on membrane capacitance, thus confirming involvement of presynaptic Ca^2+^-dependent mechanism of vesicle release. **(G)** Quantitative PCR showing the relative mRNA expression of SNCA in mDA neurons at 3 weeks of differentiation in A53T and SNCA x3 patient lines normalised to the expression in control lines (n = 3 control lines, 2 A53T lines, 1 SNCA x3 line, 1 neuronal induction, ns p > 0.05, ** p < 0.005, one-way ANOVA). Values plotted as ±SEM. **(H)** Representative immunocytochemistry images showing the expression of alpha-synuclein in control, A53T, and SNCA x3 patient mDA neurons at 4 weeks of differentiation (scale bar = 50μm). **(I)** Quantification of the relative normalised fluorescence intensity of alpha-synuclein in control, A53T, and SNCA x3 patient mDA neurons at 4 weeks of differentiation (n = 2 control lines, 2 A53T lines, 1 SNCA x3 line, 1 neuronal induction, 3-6 fields of view per line, ns p > 0.05, * p < 0.05, ** p < 0.005, one-way ANOVA). Values plotted as ±SEM. **(J)** Representative immunocytochemistry images showing the expression of filamentous aggregated forms of alpha-synuclein recognised by a conformation specific antibody, in mDA neurons at 6 weeks of differentiation (scale bar = 10μm). **(K)** Quantification of the normalised fluorescence intensity of aggregated filamentous forms of alpha-synuclein from control, A53T, SNCA x3 lines at 6 weeks of differentiation (n = 3 control lines, 2 A53T lines, 1 SNCA x3 line, 1 neuronal induction, 3-7 fields of view per line, ** p < 0.005, one-way ANOVA). Values plotted as ±SEM. **(L)** Quantification of the average puncta size of the aggregated filamentous forms of alpha-synuclein from control, A53T, SNCA x3 lines at 6 weeks of differentiation (* p < 0.05, ** p < 0.005, one-way ANOVA). Values plotted as ±SEM.

### Patient-specific mDA neurons exhibit characteristic alpha-synuclein proteinopathy

Using RT-qPCR, we demonstrate a 4-fold increase in SNCA mRNA in the SNCA x3 line compared to the control lines, at 3 weeks post differentiation (Fig. 4G). Interestingly, the A53T lines exhibited a similar amount of SNCA mRNA compared to the control lines (Control = 1.0 ± 0.19, A53T = 1.4 ± 0.02, SNCA x3 = 4.6, p < 0.005). Using an antibody that recognises total alpha-synuclein, we detected alpha-synuclein protein expression across all lines at 4 weeks post differentiation, with a significant observable increase in the expression of alpha-synuclein in the SNCA x3 line (Fig. 4H, Fig. 4I) (Control = 1.00 ± 0.03, A53T = 1.05 ± 0.02, SNCA x3 = 1.24 ± 0.06, p < 0.005). During disease, alpha-synuclein aggregates within neurons. To test whether the SNCA mDA neurons exhibit aggregate formation, we utilised a conformation specific antibody that recognises all types of aggregated forms of alpha-synuclein (Angelova *et al*., 2020). At 6 weeks post differentiation, we observed the presence of aggregated alpha-synuclein in both the A53T and the SNCA x3 line which were significantly increased compared to control (Fig. 4J), based on fluorescence intensity (Fig. 4K) (Control = 1.00 ± 0.04, A53T = 1.21 ± 0.05, SNCA x3 = 1.18 ± 0.03, p < 0.005), and the average size of the aggregate puncta (Fig. 4L) (Control = 0.52μm^2^ ± 0.06, A53T = 0.94μm^2^ ± 0.11, SNCA x3 = 0.99μm^2^ ± 0.11, p < 0.05; p < 0.005).

### mDA neurons carrying SNCA mutations exhibit oxidative stress and mitochondrial and lysosomal dysfunction

Lastly, we used the mDA neurons from patients to probe for molecular PD phenotypes. We have previously shown that hiPSC derived neurons from PD patients carrying mutations linked to *SNCA* display oxidative stress and mitochondrial dysfunction (Little *et al*., 2018; Ludtmann *et al*., 2018) in cortical neurons at approximately day 70-80 in culture. We therefore attempted to verify whether mDA neurons induced with our protocol show similar phenotypes, but at earlier time points. At week 4 post differentiation (day 48 from hiPSC), we measured the generation of superoxide using the fluorescent reporter dihydroethidium (HEt) which changes fluorescence emission upon oxidation. We demonstrate the rate of superoxide production is significantly higher in the A53T lines, and approximately 40% higher in the SNCA x3 line compared to controls (Fig. 5A, Fig. 5D) (Control = 100% ± 4.9, A53T = 172.6% ± 17.2, SNCA x3 = 141.9% ± 15.4, p < 0.05) (Supplementary Fig. 4A). Next, we measured the levels of the endogenous antioxidant glutathione using the MCB indicator, and showed a significant reduction of glutathione in the A53T lines, and a non-significant reduction of glutathione in the SNCA x3 line compared to the controls (Fig. 5B, Fig. 5E) (Control = 100% ± 3.0, A53T = 78.8% ± 2.6, SNCA x3 = 91.5% ± 4.4, p < 0.0001) (Supplementary Fig. 4C). The combination of increased ROS production together with depleted antioxidant levels suggests the A53T mDA neurons are under oxidative stress. Interestingly, SNCA x3 mDA neurons did not display similar levels of oxidative stress, but appeared to display milder oxidative stress phenotypes.

**Figure 5.**
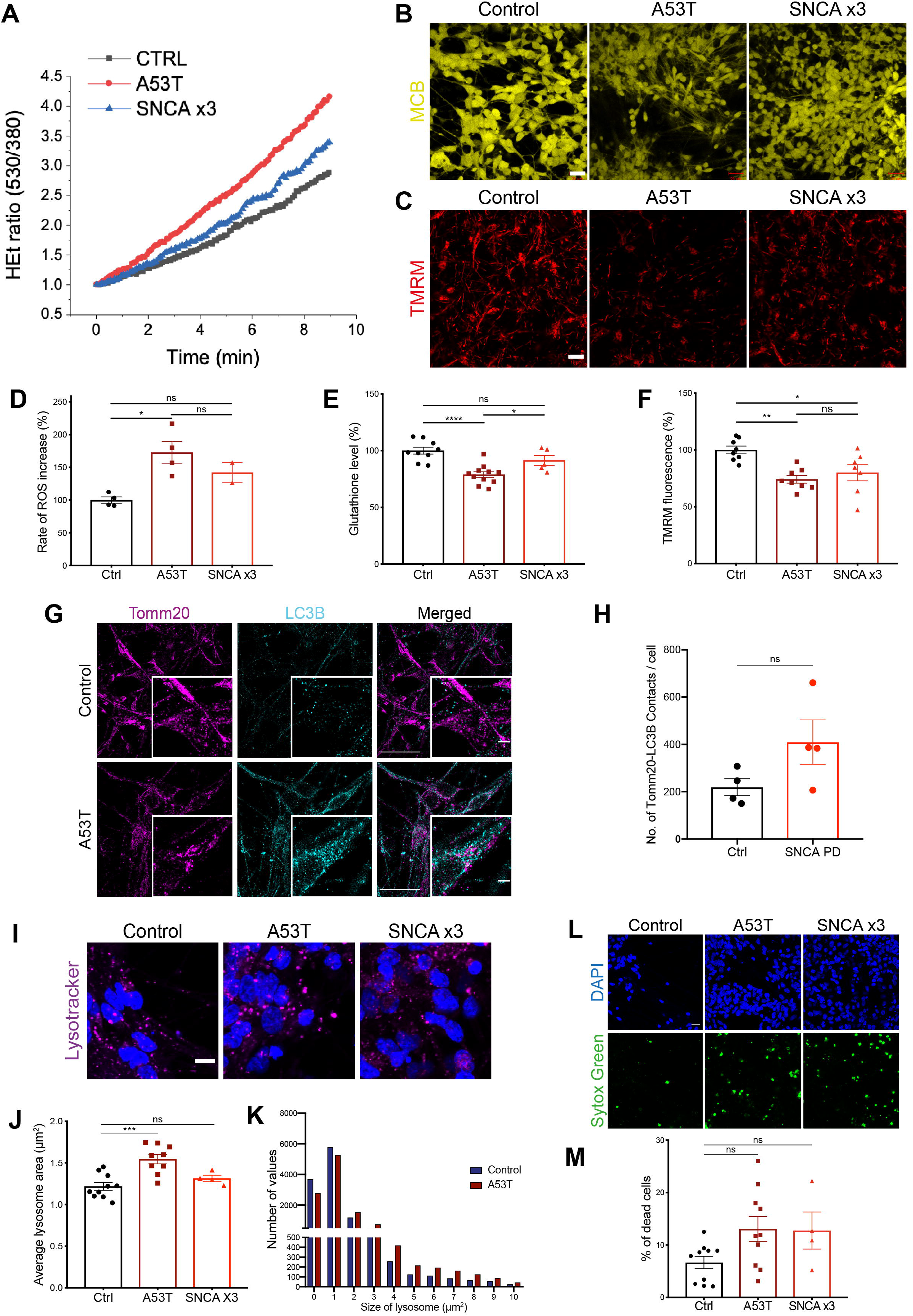
Modeling familial PD in enriched mDA neuronal cultures. **(A)** Trace showing the ratiometric measurements of superoxide generation using the fluorescent reporter dihydroethidium (HEt) in control, A53T, and SNCA x3 lines at 4 weeks of differentiation. **(B)** Representative live cell imaging of endogenous glutathione using the fluorescent reporter MCB in control, A53T, and SNCA x3 lines at 4 weeks of differentiation (scale bar = 20μm). **(C)** Representative live cell imaging of mitochondrial fluorescence using the lipophilic cationic dye TMRM in control, A53T, and SNCA x3 lines at 4 weeks of differentiation (scale bar = 10μm). **(D)** Quantification of the rate of superoxide generation based on HEt ratiometric fluorescence (n = 2 coverslips per line, across 2 control lines, 2 A53T lines, and 1 SNCA x3 line, 1 neuronal induction, ns p > 0.05, * p < 0.05, one-way ANOVA). Values plotted as ±SEM. **(E)** Quantification of the endogenous level of glutathione based on MCB fluorescence (n = 4-6 fields of view per line; 2 coverslips per line, 2 control lines, 2 A53T lines, and 1 SNCA x3 line, 1 neuronal induction ns p > 0.05, * p < 0.05, **** p < 0.0001, one-way ANOVA). Values plotted as ±SEM. **(F)** Quantification of the normalised fluorescence intensity of the mitochondrial dye TMRM (n = 3-4 fields of view per line, 2 coverslips per line, across 2 control lines, 2 A53T lines, and 1 SNCA x3 line, 2 neuronal inductions, ns p > 0.05, * p < 0.05, ** p < 0.005, one-way ANOVA). Values plotted as ±SEM. **(G)** Structured illumination microscopy (SIM) images of control and A53T cells probed for mitochondrial marker Tomm20, and the autophagosome marker LC3B, in 5 week terminally differentiated mDA neurons (scale bar = 20μm). Smaller image depicts a zoomed in version showing morphology and colocalization of the markers (scale bar = 2μm). **(H)** Quantification of the number of Tomm20-LC3B colocalizations per cell in control and familial SNCA PD lines at 5 weeks of differentiation (n = 1-2 images per line, 2 control lines, 2 A53T lines, 1 SNCA x3 line, 1 neuronal induction, ns p > 0.05 (p = 0.11), unpaired t-test). Values plotted as ±SEM. **(I)** Representative live cell imaging of lysosomes and the nuclear marker Hoechst 33342 form control, A53T, and SNCA x3 mDA neurons at 4 weeks of neuronal differentiation (scale bar = 5μm). **(J)** Quantification of the average lysosomal area from control, A53T, and SNCA x3 lines after 4 weeks of differentiation (n = 4-6 images per line, across 2 control lines, 2 A53T lines, and 1 SNCA x3 line, 1 neuronal induction ns p > 0.05, *** p < 0.0005, one-way ANOVA). Values plotted as ±SEM. **(K)** Histogram plot showing the relative number of lysosomes in each set area bin (0-10μm^2^) in control and A53T lines after 4 weeks of differentiation (n = 3-5 images per line, across 2 control lines, 2 A53T lines, 1 neuronal induction). **(L)** Live cell images depicting dead cells in mDA neurons at 4 weeks of terminal differentiation using the fluorescent dye SYTOX green in control, A53T, and SNCA x3 lines (scale bar = 20μm). **(M)** Quantification of the percentage of dead cells based on SYTOX green imaging in control and familial SNCA PD lines (n = 4-5 fields of view, 2 control lines, 2 A53T lines, and 1 SNCA x3 line, 1 neuronal induction, ns p > 0.05, one-way ANOVA). Values plotted as ±SEM.

We next assessed mitochondrial function at 4 weeks using the lipophilic cationic dye TMRM, which accumulates less in depolarised mitochondria resulting in lower TMRM signal upon the loss of mitochondrial membrane potential. We showed a reduction in mitochondrial membrane potential in both the A53T and SNCA x3 lines compared to the control neurons, suggesting potential mitochondrial dysfunction (Fig. 5C, Fig. 5F) (Control = 100% ± 3.4, A53T = 74.1% ± 3.3, SNCA x3 = 80.0% ± 7.1, p < 0.05, p < 0.005) (Supplementary Fig. 4B).

We also investigated whether lysosomal dynamics were affected at 4 weeks post differentiation. Mature mDA neurons were loaded with the lysosomal dye LysoTracker Deep Red and imaged to visualise lysosomes (Fig. 5I). We found that A53T neurons, but not SNCA x3 neurons exhibited significantly larger lysosomes compared to control samples (Fig. 5J) (Control = 1.2μm^2^ ± 0.05, A53T = 1.5μm^2^ ± 0.06, SNCA x3 = 1.3μm^2^ ± 0.04, p < 0.0005). Similarly, by plotting the area of lysosomes in a histogram, we observed a shift in lysosomal area in A53T samples compared to controls. Control samples had a higher proportion of smaller lysosomes (mainly between 0-2μm^2^) compared to the A53T neurons, which had a higher proportion of larger lysosomes (mostly 2-10μm^2^) (Fig. 5K), suggesting potential lysosomal dysfunction. Both damaged mitochondria as well as protein aggregates are cleared through the autophagy pathway in lysosomes. Therefore, we studied the expression of the autophagosome marker LC3B together with the mitochondrial marker Tomm20 at week 5 post differentiation (day 55). We adopted a super resolution approach, structured illumination microscopy (SIM) to resolve the contacts between the mitochondria and the autophagosomes. Control cells exhibited Tomm20 staining with an intact mitochondrial network and exhibited a low level of LC3B staining (top panels, Fig. 5G). In contrast, the Tomm20 appearance in the A53T lines exhibited fragmented mitochondria with a disrupted mitochondrial network, and there was an increase in the LC3B staining suggesting an autophagosome response (lower panels, Fig. 5G). We also additionally observed an increase in the number of colocalizations between Tomm20 and LC3B in all SNCA PD lines compared to control lines, suggesting close contact between autophagosomes and the mitochondria (Fig. 5H) (Control = 219 ± 36, SNCA PD = 410 ± 94, Tomm20-LC3B colocalizations per cell, p > 0.05).

Finally, we examined the basal viability of the SNCA mutant neurons compared to the controls using the fluorescent dye SYTOX-green to identify dead cells. We observed an increase in cell death of above 10% in both A53T and SNCA x3 lines compared to control lines at 4 weeks post differentiation (Fig. 5L, Fig. 5M) (Control = 6.7% ± 1.2, A53T = 13.1% ± 2.4, SNCA x3 = 12.8% ± 3.5, p > 0.05) (Supplementary Fig. 4D). Additionally, pyknotic cells were also higher in SNCA PD lines compared to control lines (Control = 14.9% ± 1.3, SNCA PD = 19.3% ± 1.8, p > 0.05) (Supplementary Fig. 4E, 4F). In summary, we have generated highly enriched cultures of mDA neurons from patient hiPSCs harbouring mutations that cause PD, and demonstrate that these are functionally mature, and can be utilised to investigate the pathological role of synuclein.

## Discussion

Lineage restriction to diverse cellular fates in the neuraxis is the consequence of interplay of multiple developmental signals, which are regulated in a spatio-temporal manner. The development of protocols based on these developmental insights to successfully differentiate hiPSCs into enriched mDA neurons is vitally important for modelling PD, as well as for cell replacement therapies. Here we successfully generated enriched mDA NPCs and neurons using only small molecules. We initiated neuronal induction via small molecule dual SMAD inhibition (Chambers *et al*., 2009), caudalization to the midbrain via timed activation of Wnt signalling through a GSK-3 beta inhibitor, and floor-plate ventralization through Shh signalling activated by small molecule agonism. Applying developmentally rationalized patterning cues to hiPSCs with our protocol yielded enriched mDA progenitors, which were subsequently differentiated into enriched mDA TH+ neurons (>80%) after 3 weeks (41 days from the pluripotent state). Single-cell RNA-seq identified 7 neuronal clusters, which expressed key markers of mDA neurons, and a further 5 clusters with the identity of midbrain NPCs. This confirmed that the culture is representative of the developing midbrain, with the appearance of abundant mDA neurons at week 4 of differentiation (48 days from pluripotent state), similar to previous studies using human fetal midbrain tissue (La Manno *et al*., 2016). Our mDA neurons displayed functional properties including calcium channel activity as early as 2 weeks of differentiation, and to our knowledge, the first time the fluorogenic DAT dye was used in hiPSC systems to show expression, and functionality of the DAT. We also show our cultures display various electrophysiological characteristics, which go on to form functional neuronal networks as they age. Finally, we show our mDA cultures can successfully model various common PD phenotypes.

Though protocols have been developed to produce enriched ventral midbrain NPCs and mDA neurons (Kirkeby *et al*., 2017; Kriks *et al*., 2011), these protocols are often expensive due to the use of neurotrophic growth factors, and are variable when using hiPSCs for modelling PD. The use of these growth factors and recombinant proteins to pattern cells to the correct part of the ventral midbrain, to yield enriched progenitors has been shown to be vitally important in graft outcome (Kee *et al*., 2017; Kirkeby *et al*., 2017). However, we interestingly found that our mDA NPCs were still able to successfully differentiate into enriched mDA neurons without the use of any neurotrophic growth factors, which have been shown to be important for the maintenance and survival of mDA neurons (Peng *et al*., 2011). We used a ROCK inhibitor to enhance cell survival, and a Notch inhibitor to promote cell cycle exit. Using a combination of these two molecules allowed our mDA NPCs to survive and exit the cell cycle, promoting differentiation into post-mitotic mDA neurons. The enrichment and cost effectiveness underlie the usefulness of our protocol to model PD over current protocols.

Differentiating hiPSCs from patients with PD also yielded highly enriched mDA neurons, which displayed electrophysiological characteristics. Oxidative stress has been reported in synucleinopathy, and in hiPSC derived cortical neurons carrying the SNCA x3, where we observed an increase in superoxide and hydrogen peroxide at day 80 (Deas *et al*., 2016). Interestingly on differentiation to mDA neurons, we observed aberrant generation of superoxide, with concomitant reduction in glutathione at day 28. This suggests that the more vulnerable neuronal population in PD exhibits cellular pathology at an earlier stage in vitro. Similarly, In previous studies mitochondrial dysfunction in A53T, and SNCA x3 hiPSC-derived cortical neurons arose between days 50-80 (Chung *et al*., 2013; Ludtmann *et al*., 2018). In mDA neurons, mitochondrial depolarisation was detected between 50-60 days (Little *et al*., 2018). Using our protocol, we detected mitochondrial dysfunction in the form of mitochondrial depolarisation at day 48 post differentiation, which could be due to the enhanced vulnerability of cells due to a higher enrichment of mDA neurons. This will allow the unmasking of early molecular phenotypes, in comparison to cortical neurons. At a similar age, we detected an alteration in the lysosomes in the A53T lines, and impaired lysosomal capacity has been previously reported (Mazzulli, Zunke, Isacson, *et al*., 2016). Furthermore, super resolution microscopy revealed mitochondrial fragmentation, in keeping with the detection of mitochondrial dysfunction. There was also an increase in autophagosomes, and the autophagic response is in close physical contact with fragmented mitochondria, similar to previous studies (Ryan *et al*., 2018). Our results support a hypothesis that both the lysosomal response to alpha-synuclein and the mitochondrial response to alpha-synuclein are altered in SNCA iPSC models, and that concomitant impaired removal of misfiled protein and damaged mitochondria may lead to neuronal death.

Here we used the conformation-specific filament antibody which allowed the aggregates to be distinguished from the total alpha-synuclein population. We were able to detect the presence of aggregated forms of alpha-synuclein, which were higher in all SNCA PD lines, at week 6 of differentiation (day 62), which is earlier than previously reported (Mazzulli, Zunke, Tsunemi, *et al*., 2016). Though aggregates have been reported as early as day 35 in mDA cultures (Zambon *et al*., 2019), aggregation is notoriously hard to detect in cell model systems. Using single molecule and super resolution approaches (Ludtmann *et al*., 2018), it may be possible to resolve the aggregation process at higher resolution, and detect the formation of small soluble oligomers within cells at earlier time points.

In summary, we successfully generate highly enriched population of mDA neurons from hiPSCs, that express mDA markers, functional dopamine transport and form neuronal networks. Our approach is more efficient, and economically favourable than other protocols. We were able to use this approach to generate a PD model based on SNCA mutations. We were able to take a highly vulnerable population of neurons in PD and determine the temporal sequence of events as they arise in culture. At 4 weeks of differentiation, oxidative stress, mitochondrial dysfunction, and lysosomal responses were seen in cultures. As the neurons aged, we observed an increase in autophagic abnormalities, and fragmentation of the mitochondria at week 5. Finally, we were able to observe an increase in alpha-synuclein aggregates by week 6 (Fig. 6). Through the generation of the most vulnerable neurons in Parkinson’s disease, our model recapitulates all the hallmarks of synucleinopathy at earlier time points in culture, concomitant with neuronal maturation. Our approach allows these established hallmarks, and their emergence to be temporally sequenced in our model, and critically allows new disease mechanisms to be identified at the earliest possible stages.

**Figure 6.**
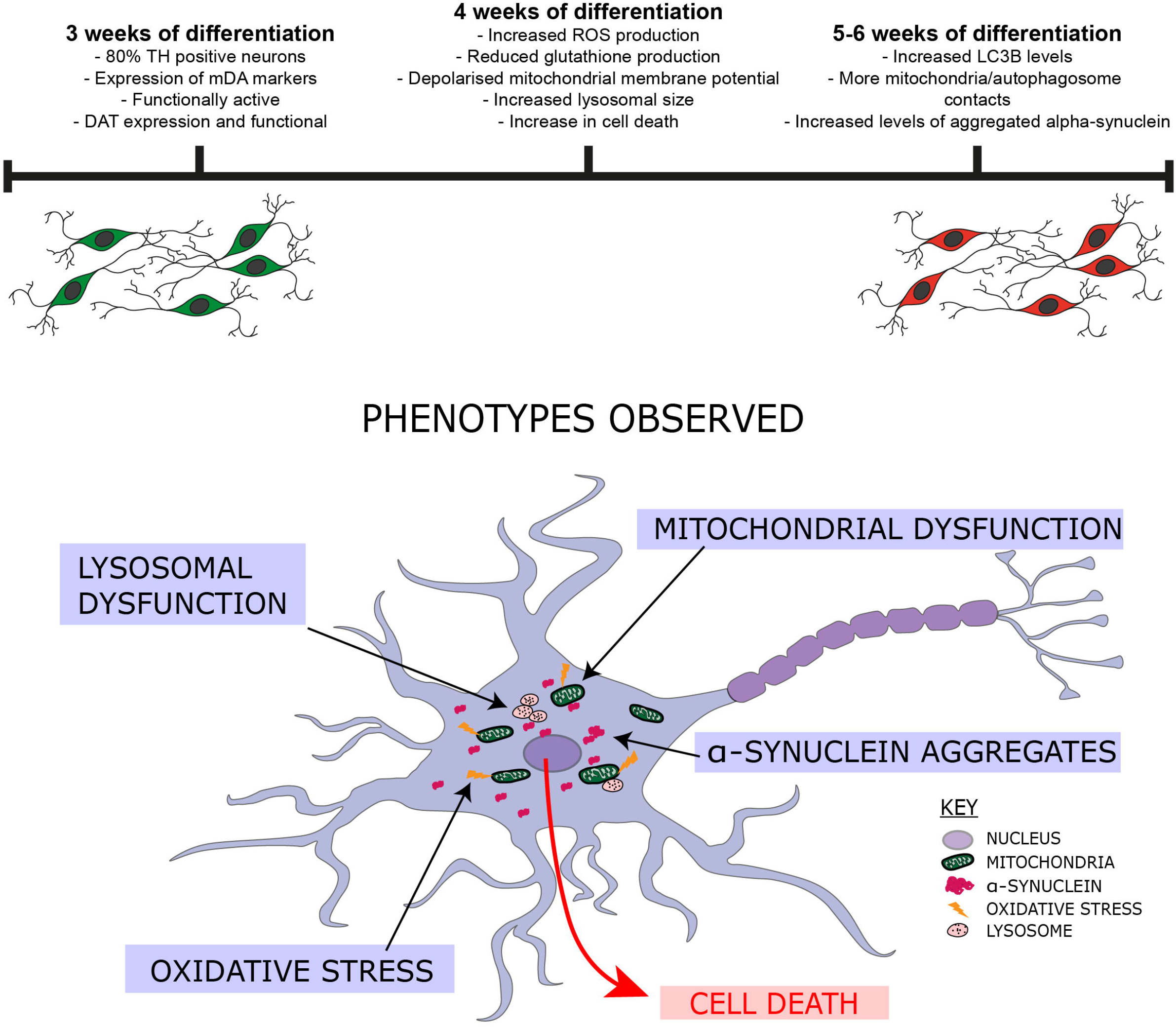
Highly enriched mDA neurons model common phenotypes associated with synucleinopathy. The top illustration shows that our protocol yields highly enriched (80%) mDA neurons after 3 weeks of differentiation. Patient derived neurons go on to develop oxidative stress, mitochondrial and lysosomal dysfunction after 4 weeks. After 5-6 weeks there is an impairment in mitophagy, as well as aggregation of alpha-synuclein. The bottom illustration highlights the phenotypes we detected in this study using our mDA neuron differentiation protocol.

## Abbreviations

DAT: Dopamine transporter
FGF: Fibroblast growth factor
GDNF: Glial cell-derived neurotrophic factor
GIRK2: G protein-activated inward rectifier potassium channel 2
hiPSC: Human induced pluripotent stem cells
mDA: Midbrain dopaminergic
NPC: Neuronal precursor cell
Nurr1: Nuclear receptor related-1 protein
PD: Parkinson’s disease
ROCK: Rho-associated protein kinase
ROS: Reactive oxygen species
SHH: Sonic hedgehog
SIM: Structured illumination microscopy
SNCA: Alpha-synuclein gene
TH: Tyrosine hydroxylase
Wnt1: Wingless-int1

## Acknowledgements

We would wish to thank the patients for the fibroblast donation. We would also like to thank the Francis Crick Institute Flow Cytometry, Advanced Sequencing, and Bioinformatics and Biostatistics STPs for their help in conducting and analysing the flow cytometry, and single-cell RNA-seq experiments.

## Supplementary material

### Supplementary methods

#### Midbrain dopaminergic neuron (mDA) differentiation

For all mDA differentiations, hiPSCs were grown to 100% confluency. Differentiation was triggered by removing old media and replacing it with a chemically defined medium consisting of DMEM/F12, N2 supplement, Neurobasal, B27 supplement, L-Glutamine, non-essential amino acids, 50U/ml penicillin-streptomycin, β-mercaptoethanol (all from ThermoFisherScientific) and 5μg/ml insulin (Sigma), termed “N2B27”. Cells were patterned for 14 days with daily media changes. For the first 2 days, the media was supplemented with the small molecules 5μM SB431542 (Tocris Bioscience), 2μM Dorsomorphin (Tocris Bioscience), 1μM CHIR99021 (Miltenyi Biotec). On day 2, 1μM Purmorphamine (Merck Millipore) was added. On day 8, CHIR99021, and SB431542 were removed leaving only Dorsomorphin and Purmorphamine in the medium until day 14. Cells were enzymatically dissociated and split on days 4, 10, and 14 using 1mg/ml Dispase (ThermoFisherScientific). After patterning, mDA neuronal precursor cells (NPCs) were maintained in N2B27 for 4 days. On day 19, cells were plated onto Geltrex pre-coated Ibidi 8-well chambers (100k/well), clear bottom 96-well plates (50k/well), or 12-well plates (500k/well), and terminally differentiated using N2B27 supplemented with 0.1μM Compound E (Enzo Life Sciences) and 10μM Y-27632 dihydrochloride (Rho kinase ROCK inhibitor) (Tocris) from day 20 for the whole duration of terminal differentiation, with two weekly media changes. NPCs could be cryopreserved on day 19, before terminal differentiation. For super-resolution microscopy, cells were plated on glass-bottom Ibidi 8-well chambers which were pre-coated with poly-D-lysine (PDL) overnight, followed by laminin (Sigma, L2020) in PBS for 1 hour.

#### Immunocytochemistry (ICC)

For ICC, cells at the desired time point of differentiation had the media removed, followed by one wash in PBS, and fixed with 4% paraformaldehyde for 15 minutes at room temperature (RT). The paraformaldehyde was then removed, and cells were washed once in PBS. For the LC3B antibody (Cell Signalling Technologies #3868), samples were incubated in -20°C methanol after paraformaldehyde fixation. All samples were then blocked for non-specific binding and permeabilized in 5% bovine serum albumin (BSA) (Sigma) + 0.2% Triton X-100 (Sigma) in PBS for 60 minutes. The primary antibodies were then made up to the desired dilution (Table 1) in 5% BSA and applied to the cells overnight at 4°C. The cells were then washed twice in PBS followed by the application of species-specific, secondary antibodies conjugated to relative Alexa Fluor dyes (ThermoFisherScientific), at a 1:500 dilution made up in 5% BSA to cells for 60 minutes at RT in the dark. After secondary antibody incubation, cells were washed once in PBS before being stained with 4′,6[diamidino[2[phenylindole nuclear stain (DAPI) in PBS for 5 minutes at a 1:1000 dilution. After DAPI incubation, cells were washed once with PBS before being submerged in fluorescence mounting medium (Dako), and stored at 4°C until imaging.

The samples were imaged using Zeiss 880 confocal system with a 40x, 1.4 N.A. oil objective, and a pinhole of 1 airy units (AU). Between 3-5 images were collected per sample, all with a Z projection consisting of 5 slices, and displayed as a maximum projection. Samples were also imaged using the PerkinElmer Opera Phenix™High Content Screening System with 20 and 40x water objective lenses. A minimum of 5 fields of view and a Z projection of 3 slices was acquired per well, with the images displayed as a maximum projection. The accompanying software, Columbus™was used to store and analyse acquired images. The settings for the acquisition of images were kept the same for all samples in the experiment set.

#### RNA extraction and quantitative polymerase chain reaction (qPCR)

RNA was harvested from snap frozen cell pellets using the Maxwell®[RSC simplyRNA Cells kit (Promega), and the accompanying Maxwell®[RSC 48 instrument. After RNA extraction, the RNA concentration and quality using the 260/280 ratio was assessed using the nanodrop. Up to 1μg of RNA was retro-transcribed into cDNA using the High-Capacity cDNA Reverse Transcription kit (ThermoFisherScientific). The qPCR was performed using TaqMan™Gene Expression Assay (ThermoFisherScientific). For each gene, TaqMan™probes were used (Table 2) along with the TaqMan™master mix, and sample cDNA following the manufacturer’s protocol. Samples, along with a minus reverse transcriptase control (-RT) were ran for each gene on the QuantSudio 6 Flex Real-Time PCR System (Applied Biosystems). The -RT served as a negative control, and the gene expression levels were normalised to the housekeeping gene GAPDH following the delta-delta Ct method. Gene expression values were expressed as the normalisation to either hiPSCs or mDA NPCs.

### Single-cell RNA-seq

#### Single cell generation, cDNA synthesis, library construction, and sequencing protocol

After 4 weeks of differentiation, mDA neurons from 3 control hiPSC lines were washed with PBS once, and then incubated with Accutase (Gibco™) for 5 minutes to obtain a single cell suspension. The samples were then diluted 1/3 before usage. The quality and concentration of each single-cell suspension was measured using Trypan blue and the Eve automatic cell counter. Each sample presented a concentration between a 1200-1700 cell/µl and viability ranged between 55-68%, samples with a viability above 57% were used for sequencing. Approximately 10000 cells were loaded for each sample into a separate channel of a Chromium Chip G for use in the 10X Chromium Controller (cat: PN-1000120). The cells were partitioned into nanoliter scale Gel Beads in emulsions (GEMs) and lysed using the 10x Genomics Single Cell 3′ Chip V3.1 GEM, Library and Gel Bead Kit (cat: PN-1000121). cDNA synthesis and library construction were performed as per the manufacturer’s instructions. The RNA was reversed transcribed and amplified using 12 cycles of PCR. Libraries were prepared from 10µl of the cDNA and 13 cycles of amplification. Each library was prepared using Single Index Kit T Set A (cat: PN-1000213) and sequenced on the HiSeq4000 system (Illumina) using 100 bp paired-end run at a depth of 65-100 million reads. Libraries were generated in independent runs for the different samples.

#### Pre-processing single-cell RNA-seq (scRNA-seq) data

Using the Cell Ranger v3.0.2 Single-Cell Software Suite from 10X Genomics reads were aligned to the human reference genome (Ensembl release 93, GRCh38). The analysis was carried out using Seurat v3.1.0 (Butler *et al*., 2018; Stuart *et al*., 2019) in R-3.6.1 (R Core Team, 2019). Cells expressing fewer than 200 genes were excluded from the subsequent analysis. Using default parameter within Seurat v3.1.0 (Butler *et al*., 2018; Stuart *et al*., 2019) data for each sample were normalised across cells using the ‘LogNormalize’ function with a scale factor of 10,000. A set of highly variable genes was identified using the ‘FindVariableFeatures()’ function (selection.method = “vst”, nfeatures = 2000). Data were centred and scaled using the ‘ScaleData()’ function with default parameters. Using the highly variable genes, PCA was performed on the scaled data and the first 30 principal components were used to create a Shared Nearest Neighbour (SNN) graph using the ‘FindNeighbors()’ function (k.param = 20). This was used to find clusters of cells showing similar expression using the ‘FindClusters()’ function across a range of clustering resolutions (0.2-1.4 in 0.2 increments). Based on the visualisation of average mitochondrial gene expression across different cluster resolutions using the R package Clustree v0.4.1 (Zappia and Oshlack, 2018) we selected a clustering resolution of 1.0 to exclude cluster with an average mitochondrial gene expression above 7.5% and, concomitantly, an average number of detected features below 1300.

#### Integration across samples

After filtering of cells/clusters based on mitochondrial gene expression and the number of detected features, we integrated the three samples using the standard workflow from the Seurat v3.1.0 package (Stuart *et al*., 2019). After data normalisation and variable feature detection in the individual samples (see above), anchors were identified using the ‘FindIntegrationAnchors()’ function and datasets were integrated with the ‘IntegrateData()’ across 50 dimensions for all detected features in the datasets. We then performed dimension reduction (PC1-45) and cluster identification at resolutions 1.4. After removal of a cluster consisting mostly of suspected doublet cells identified using DoubletFinder (McGinnis *et al*., 2019), we performed data scaling including cell cycle score regression, dimension reduction (PC1-50) and cluster identification (resolutions 0.4-1.8).

Biomarker of each cluster were identified using Seurat’s ‘FindAllMarkers()’ function using the Wilcoxon rank sum test. We limited the test to positive markers for each cluster in comparison to all remaining cells. The positive marker genes had to be detected in 25% of cells in either of the two groups, with limiting testing further to genes which show, on average, at least 0.25-fold difference (log-scale) between the two groups of cells. Cluster identity was determined using visual inspection focusing on the expression of known marker genes.

#### Flow Cytometry

The protocol for staining cells for flow cytometry analysis was adapted from a previous study (Turaç *et al*., 2013). Cells were washed once with PBS, before being detached into a single cell suspension using Accutase (Gibco™). A cell suspension of 500k/ml was prepared in media. Cells were then spun down at 200g for 5 minutes, and the supernatant was removed. Cell pellet was resuspended gently in 4ml of 4% paraformaldehyde and briefly vortexed at a low speed before being spun on a rotation spinner for 10 minutes at RT. After fixation, samples were spun down and supernatant removed. Cells were resuspended in 2ml of 0.1% BSA in PBS. After resuspension, cells were filtered through a 70μm strainer (Miltenyi Biotec) to filter out any cell clumps. Cells were then spun down again, and the supernatant was removed. Cell pellets were then resuspended in 1ml of permeabilization/blocking buffer (0.1% Triton X-100, 1% BSA, 10% normal goat serum (Sigma) in PBS), and incubated on a rotation spinner for 30 minutes at RT. After permeabilization/blocking, cells were spun down and the supernatant was removed. Cells were then resuspended in the primary antibodies (1:200) made up in 0.1% BSA in PBS, and incubated on the rotation spinner for 1 hour at RT. After primary antibody incubation, cells were spun down, supernatant removed and washed once in 0.1% BSA in PBS. They were then resuspended in the species-specific secondary antibodies (AlexaFluor 488, 647) at a dilution of 1:500 made up in 0.1% BSA in PBS and incubated in the dark on a rotation spinner for 30 minutes. After incubation, cells were spun down, supernatant removed and washed once in PBS, followed by incubation with DAPI made up in PBS for 5 minutes. The DAPI + PBS was then removed, followed by one wash in PBS, before being analysed on the flow cytometer.

The samples were run on the LSRii (BD) cell sorter. Scattering was initially used to discard debris as well as cell doublets and larger clumps. The single cell population was then gated to include DAPI positive only cells (negative control). The gating threshold for measured channels was determined using the control lacking the antibody of interest (Fluorescence minus one (FMO) control), for both channels being recorded. Once the parameters had been set, 10,000 cell events were recorded, and data was processed and analysed on FlowJo.

#### Electrophysiology

Visualized patch-clamp recordings from cell cultures were performed using an infrared differential interference contrast imaging system. The perfusion solution contained the following (in mM): 119 NaCl, 2.5 KCl, 1.3 Na_2_SO_4_, 2.5 CaCl_2_, 26.2 NaHCO_3_, 1 NaH_2_PO_4_, 2 CaCl_2_, 2 MgCl_2_, 22 glucose and was continuously bubbled with 95% O_2_ and 5% CO_2_, pH 7.4. Whole-cell recordings were performed at 32-34°C; the patch-clamp pipette resistance was 3-7 MΩ depending on particular experimental conditions. Series resistance was monitored throughout experiments using a +5 mV step command, cells with very high series resistance (above 25 MΩ) or unstable holding current were rejected. The intracellular pipette solution for voltage-clamp experiments contained (in mM): 120.5 CsCl, 10 KOH-HEPES, 2 EGTA, 8 NaCl, 5 QX-314 Br-salt, 2 Na-ATP, 0.3 Na-GTP. For current-clamp experiments, the intracellular solution contained (in mM): 126 K-gluconate, 4 NaCl, 5 HEPES, 15 glucose, 1 K_2_SO_4_×7 H_2_O, 2 BAPTA, 3 Na-ATP. pH was adjusted to 7.2 and osmolarity adjusted to 295 mOsm. To isolate response of NMDA receptors we added to a perfusion solution: 50 mM picrotoxin, 20 mM NBQX, 1 mM strychnine, 1 mM CGP-55845, 100 mM MCPG, with zero Mg^2+^. To isolate response of GABA_A_ receptors, we added 50 mM APV, 20 mM NBQX, 1 mM strychnine, 1 mM CGP-55845, 100 mM MCPG. All chemicals were purchased from Tocris Bioscience. mDA neurons were tested as a subgroup of the set of generated cultures. Their electrophysiological profile hasn’t shown any legible difference from other cell lines. See Table 4 for the pattern of effects observed in neurons derived from different stem cell lines.

#### Live-cell imaging

To measure level of antioxidant, reactive oxygen species (ROS), mitochondrial membrane potential, lysosomal dynamics, calcium uptake, and cell death, a confocal microscope (ZEISS LSM 710/880 with an integrated META detection system) which has illumination intensity limited to 0.1 -0.2 % of laser output to prevent phototoxicity was used. The antioxidant level was accessed using glutathione indicator, 50μM Monochlorobimane (mBCI, ThermoFisherScientific) was incubated for 30 minutes and measured at 420 - 550 nm excited by a 405 nm laser. To measure mitochondrial membrane potential, cells were incubated with 25nM tetramethylrhodamine methyl ester (TMRM, ThermoFisherScientific) in HBSS for 40 minutes and then imaging was acquired using Zeiss LSM 880 confocal microscope. The 560 nm laser line was used to excite and it was measured above 560 nm. Approximately 3-5 fields of view with Z projections were taken per sample. To measure lysosomal dynamics, cells were incubated with 50nM LysoTracker™Deep Red (ThermoFisherScientific), and Hoechst 33342 in HBSS for 40 minutes and then imaged using Zeiss LSM 880 confocal microscope where the 405 nm, and 647 nm laser line were used to excite Hoechst 33342 and LysoTracker™Deep Red, respectively. Approximately 4-5 fields of view with Z projections were taken per sample.Calcium uptake was assessed using the dye Fluo-4 AM (ThermoFisherScientific). Cells were incubated with 5μM Fluo-4 AM in HBSS for 40 min, followed by two HBSS (Invitrogen) washes. Live-cell imaging was performed excited by a 480 nm laser and measured at 520 nm. A time-series with 5 second intervals was performed to establish basal fluorescence before 50mM KCl was added to depolarise the membrane and measure fluorescence intensity increase, and recovery. To measure cell death, live cells were incubated with 100nM SYTOX™Green Nucleic Acid Stain (ThermoFisherScientific), and the nuclear marker Hoechst 33342 for 40 minutes in HBSS. As SYTOX™Green is impermeable to live cells, only dead cells were stained, whereas Hoechst 33342 labelled all cells. Cells were imaged using the Zeiss LSM 880 confocal microscope where the 405 nm, and the 488 nm laser line were used to excite Hoechst 33342 and SYTOX Green, respectively. Approximately 4-5 fields of view with Z projections were taken per sample. To measure ROS, (mainly superoxide) a cooled camera device (CCD) was used and data were obtained on an epifluorescence inverted microscope equipped with a 20x fluorite objective. Cells were loaded with 2μM dihydroethidium (HEt, Molecular Probe) in HBSS. We generated ratios of the oxidised form (ethidium) exited at 530 nm and measured using a 560 nm longpass filter versus the reduced form with excitation at 380 nm measured at 415 - 470 nm.

#### Fluorescent false neurotransmitter (FFN) live-cell DAT imaging

To measure the presence and activity of the DAT, we utilized the commercially available fluorescent DAT and VMAT2 substrate FFN102 (Abcam, ab120866). To measure the uptake of the FFN102 dye, a field of view was first found using the brightfield settings on a Zeiss LSM 880 confocal microscope. The cells then had 10μM of the dye in HBSS added and a time-series with an exposure every 5 seconds using the 405 nm laser was started, to measure uptake of the dye into the cells. As a control to confirm specificity, samples in a different well were pre-treated with 5μM of the DAT inhibitor nomifensine (Sigma) for 10 minutes in HBSS. Nomifensine was kept in cell solution also after FFN102 was added. Once the dye had entered cells, the cells were depolarised by the addition of 50mM KCl to observe FFN102 dynamics. Approximately 20 cells were measured per condition/sample and their rate of fluorescence intensity increase was plotted.

#### Structured illumination microscopy (SIM)

Samples for SIM were cultured on glass bottom Ibidi 8-well chambers coated with laminin. They were fixed and stained for intracellular markers as described in the ICC section. SIM was performed on an Elyra PS.1 microscope (Zeiss), using a 40x oil objective (EC Plan-Neofluar 40x/1.30 Oil DIC M27). Images were acquired as 15x 0.1 µm Z-planes on a pco.edge sCMOS camera, using 5 grid rotations with the 405 nm (23µm grating period), the 488 nm (28µm grating period) and the 561 nm (34µm grating period) lasers. Images were processed and channels aligned using the automatic settings on the ZEN Black software (Zeiss).

### Image analysis

ICC images, lysosomal dynamics, and cell death were analysed using Fiji ImageJ. Live-cell imaging quantification was analysed using the ZEN software by getting the fluorescence intensities of the region of interest for each frame. All graphs and traces were plotted on Prism 8 (GraphPad), with the exception of Fluo4 traces in Figure 2A, which were analysed and plotted using Origin 2018 (Microcal Software Inc., Northampton, MA).

### Statistical analysis

Statistical analysis was performed on Prism 8. To compare two individual groups, a unpaired, two-tailed t-test was used to generate a p value. When comparing more than two individual groups, an ordinary one-way ANOVA was used with a post hoc Tukey test for multiple comparisons between groups. When comparing two individual variables, an ordinary two-way ANOVA was performed, with a correction of the False Discovery Rate for multiple comparisons between groups. A p value below 0.05 was considered to be statistically significant. Results are represented as means ± standard error of the mean (SEM), or standard deviation (SD) where stated in the figure legends. The number of hiPSC lines, number of cells, and the number of neuronal inductions used for each experiment is stated in the respective figure legend. The sizes of the sample for each experiment was selected to ensure that the technical (number of cells, numbers of fields of view, and number of coverslips), and biological (number of hiPSC lines, and number of neuronal inductions) variation was adequately captured, and is listed in the respective figure legends.

## Supplementary figure legends

**Supplementary Table 1.**
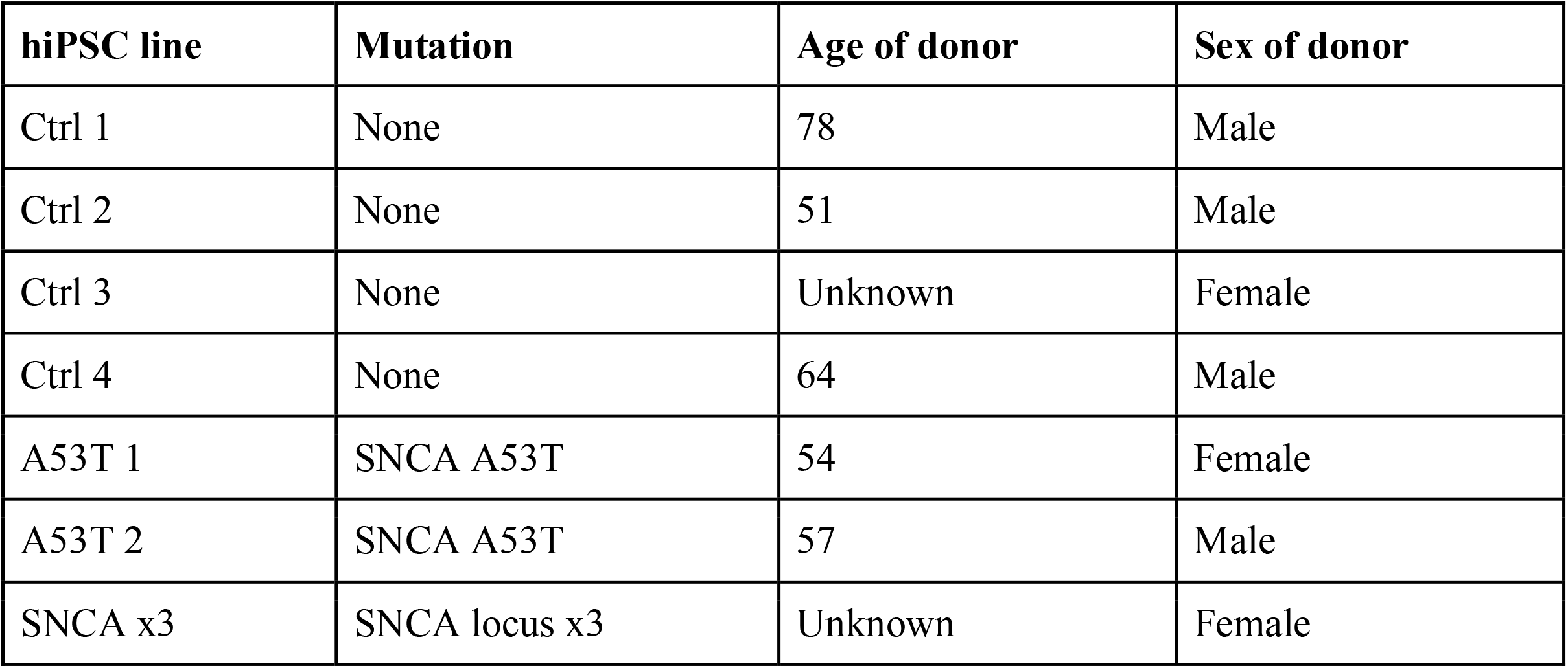
hiPSC lines used in this study.

| hiPSC line | Mutation | Age of donor | Sex of donor |
| --- | --- | --- | --- |
| Ctrl 1 | None | 78 | Male |
| Ctrl 2 | None | 51 | Male |
| Ctrl 3 | None | Unknown | Female |
| Ctrl 4 | None | 64 | Male |
| A53T 1 | SNCA A53T | 54 | Female |
| A53T 2 | SNCA A53T | 57 | Male |
| SNCA x3 | SNCA locus x3 | Unknown | Female |

**Supplementary Table 2.** List of antibodies used in this study.

| Protein | Company | Catalogue | Species | Dilution |
| --- | --- | --- | --- | --- |
| LMX1A | Abcam | ab139726 | Rabbit | 1:500 |
| FOXA2 | Santa-Cruz | sc-374376 | Mouse | 1:100 |
| OTX2 | R&D Systems | AF1979 | Goat | 1:500 |
| TH | Abcam | ab137869 | Rabbit | 1:500 (ICC)<br>1:200 (Flow) |
| TH | Abcam | ab76442 | Chicken | 1:500 |
| TUJ1 | Biolegend | 801201 | Mouse | 1:1000 (ICC) |
| Beta III Tubulin | Abcam | ab41489 | Chicken | 1:200 (Flow) |
| MAP2 | Abcam | ab11267 | Mouse | 1:500 |
| GIRK2 | Alomone Labs | APC-006 | Rabbit | 1:400 |
| Total alpha-synuclein | Abcam | ab138501 | Rabbit | 1:200 |
| Filament alpha-synuclein | Abcam | ab209538 | Rabbit | 1:50 |
| Tomm20 | Santa-Cruz | sc-17764 | Mouse | 1:1000 |
| LC3B | Cell Signaling Technologies | #3868 | Rabbit | 1:300 |

**Supplementary Table 3.** List of TaqMan™Gene Expression probes used in this study.

| **Gene** | **Catalogue Number** |
| --- | --- |
| FOXA2 | Hs00232764_m1 |
| LMX1A | Hs00898455_m1 |
| EN1 | Hs00154977_m1 |
| TH | Hs00165941_m1 |
| Nurr1 | Hs00428691_m1 |
| DAT | Hs00997374_m1 |
| GIRK2 | Hs01040524_m1 |
| SNCA | Hs00240906_m1 |
| GAPDH | Hs03929097_g1 |

**Figure S1:**
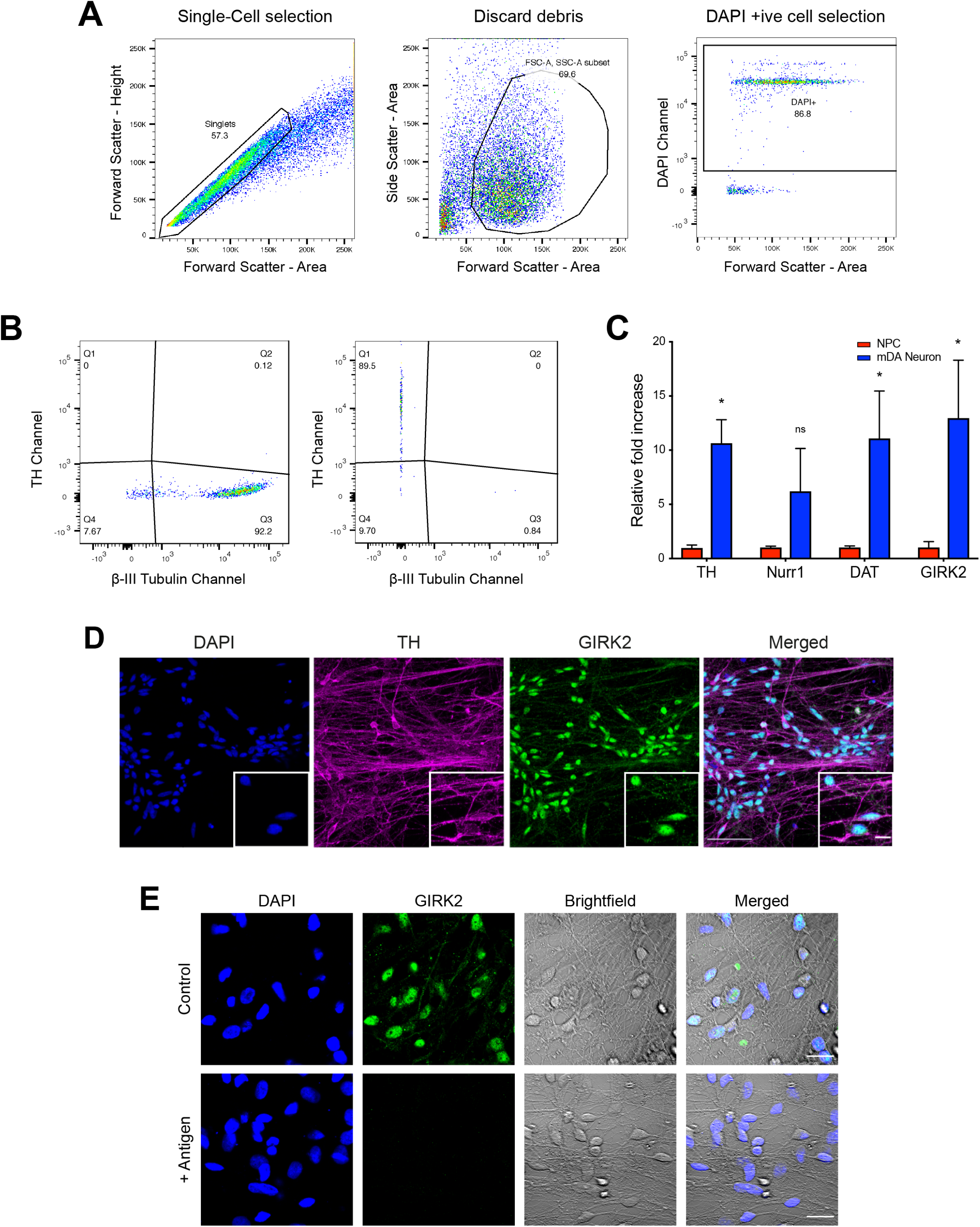
Gating threshold criteria for flow cytometry analysis and mDA neuron marker expression. **(A)** Dot plots showing the selection criteria for stained flow cytometry analysis. Using the forward scatter, only the circled cluster was selected which corresponds to single cells. Of this population, only the circled cluster was selected in the middle dot plot, discarding any debris. Finally, in the last dot plot, only the DAPI positive cells were selected, after the single cell and debris discarding gating. **(B)** Dot plots showing the thresholding gating for the channels measured. Thresholding gates were determined using the fluorescence minus one (FMO) controls. In the first dot plot, the TH gate threshold was set by measuring the intensity in the sample with all stains except the TH stain. In the next dot plot, the β-III Tubulin gate threshold was set by measuring the intensity in the sample with all stains except for the β-III Tubulin stain. **(C)** Quantitative PCR showing an up-regulation of mRNA of mature mDA markers TH, Nurr1, DAT, and GIRK2 relative to mDA NPCs (n = 3 different lines across 3 independent neuronal inductions, ns p > 0.05, * p < 0.05, ordinary two-way ANOVA). Values plotted as ±SEM. **(D)** Representative immunocytochemistry images showing a high expression of TH as well as the expression of the A9 specific mature mDA neuron marker GIRK2 (scale bar = 50μm). Smaller image depicts a zoomed in version of the image showing co-expression of TH and GIRK2 (scale bar = 10μm). **(E)** GIRK2 Antibody specificity confirmed using the control antigen. The top panel shows GIRK2 staining using the antibody as normal. The lower panel shows GIRK2 staining is abolished when the antibody was incubated with the antigen (GIRK2 sequence) the antibody was raised to, showing antibody specificity. Scale bar = 20μm.

**Figure S2:**
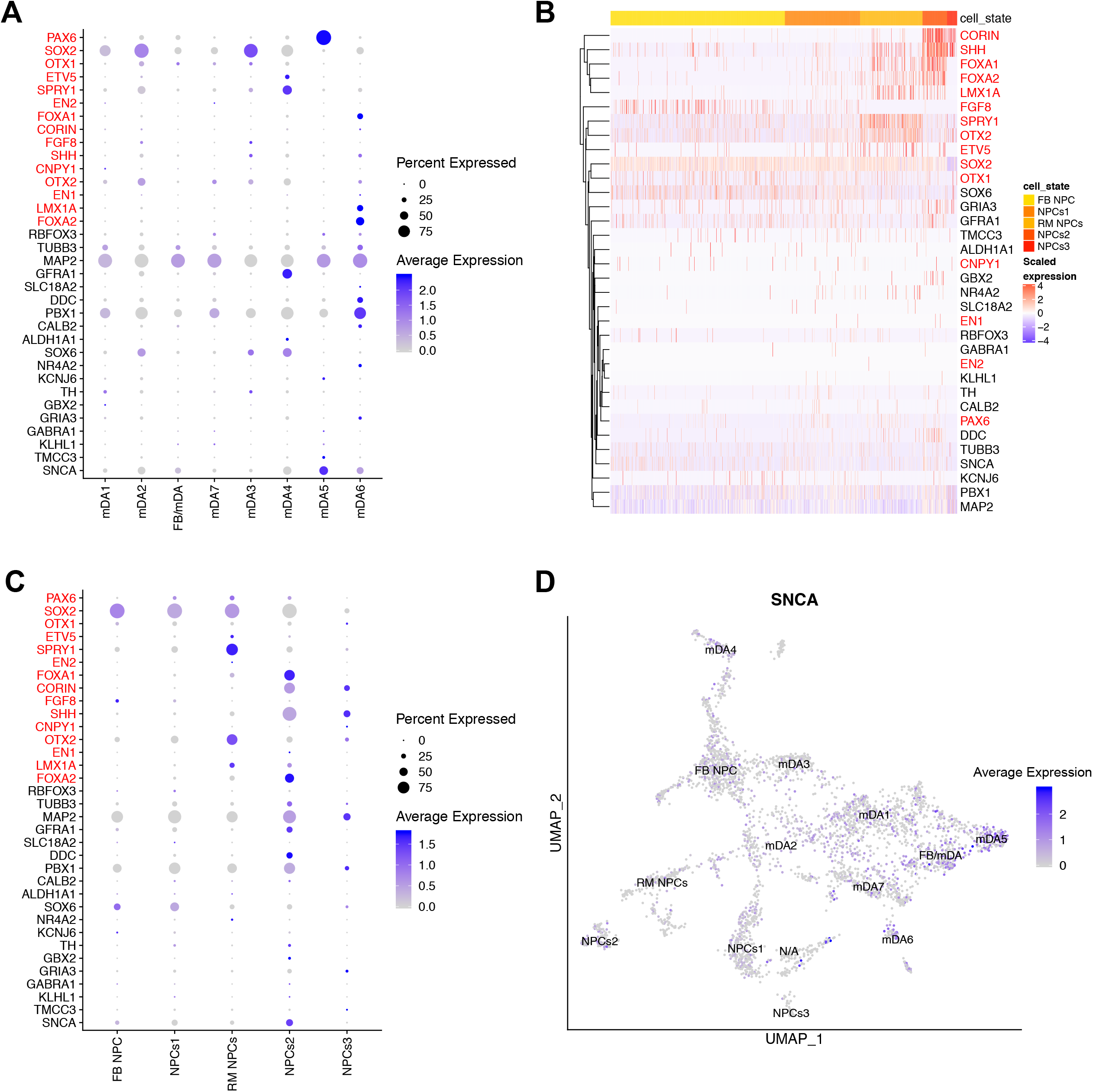
Single-cell RNA-seq analysis of mDA neurons after 4 weeks of differentiation. **(A)** A dot-plot showing the gene expression profile of the mDA clusters (mDA1-7) and the forebrain/midbrain neuron (FB/mDA) cluster identified in single-cell RNA-seq in cultures after 4 weeks of differentiation. Genes annotated in red correspond to mDA NPC markers, and those in black correspond to mDA neuron markers. Expression of the genes are coloured based on the expression fold difference and plotted based on the percentage of cells in that cluster expressing the gene. **(B)** A heat map showing the expression of genes in the clusters identified as mDA NPCs (NPCs1-3), forebrain NPCs (FB NPCs), and rostral midbrain NPCs (RM NPCs) in the culture after 4 weeks of differentiation. Each line represents a cell from that cluster. **(C)** A dot-plot showing the gene expression profile of the NPC clusters in cultures after 4 weeks of differentiation. Genes annotated in red correspond to mDA NPC markers, and those in black correspond to mDA neuron markers. Expression of the genes are coloured based on the expression fold difference and plotted based on the percentage of cells in that cluster expressing the gene. **(D)** A feature plot showing the expression of SNCA in all the clusters identified through single-cell RNA-seq at 4 weeks of differentiation.

**Figure S3:**
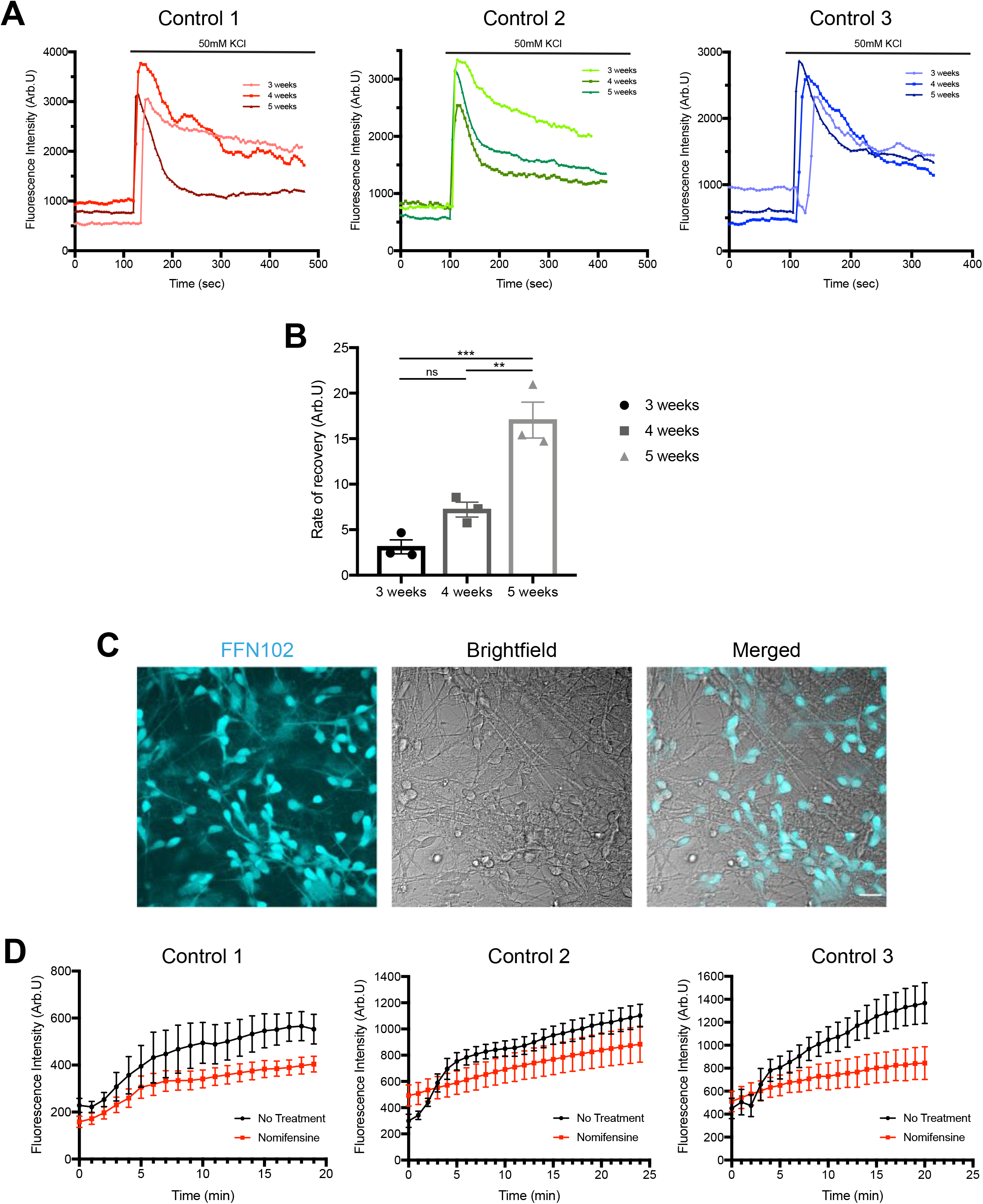
Measuring Ca^2+^ and DAT functionality in mDA neurons. **(A)** Traces showing the Fluo-4 mean fluorescence intensity for each hiPSC line before and after the addition of 50mM KCl at different weeks of differentiation (n = 15 cells per trace). Values plotted as the mean of all cells per time point of differentiation. **(B)** Quantification of the slope of fluorescence decrease after the addition of 50mM KCl for each time point, showing the Ca^2+^ efflux recovery rate increase as mDA neurons age (n = 3 hiPSC lines, ns p > 0.05, ** p < 0.005, *** p = 0.0007, one-way ANOVA). Values plotted as ±SEM. **(C)** Live-cell imaging picture of cells after a 30-minute incubation with FFN, showing the dye enters most mDA neurons. Scale bar = 20μm. **(D)** Traces showing the fluorescence intensity of FFN inside cells to measure the uptake of the dye in each hiPSC line tested, in the absence, or presence of DAT inhibitor nomifensine (n = 15-20 cells per condition). Values plotted as ±SD.

**Figure S4:**
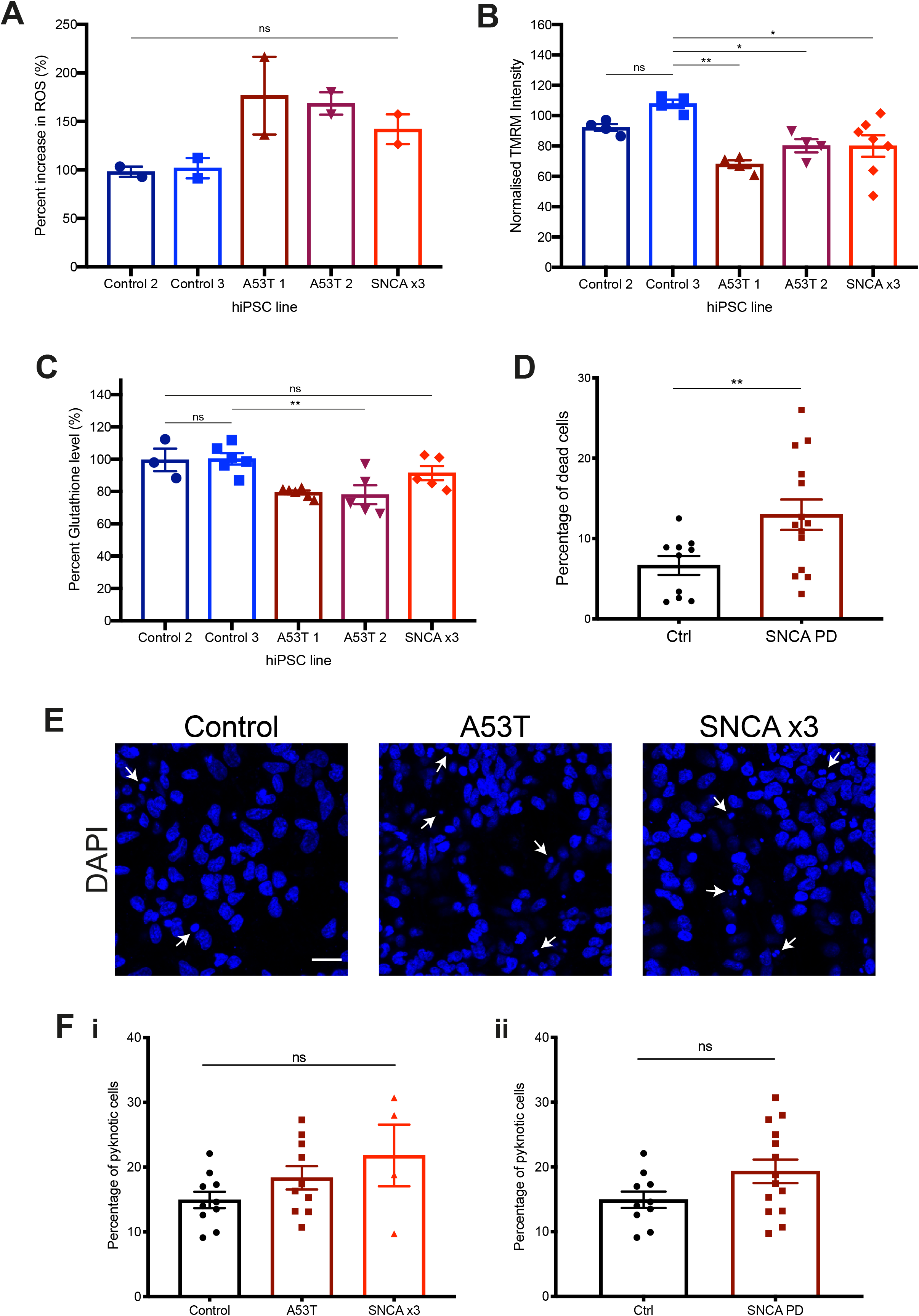
Characterisation of molecular phenotypes in SNCA PD hiPSC lines. **(A)** Relative increase in ROS based on HEt ratiometric fluorescence plotted out for each line tested (n = 2 coverslips per line, ns p > 0.05, one-way ANOVA). Values plotted as ± SEM. **(B)** The normalised TMRM fluorescence intensity plotted out for each line tested (n = 2-4 fields of view per line, across 2 coverslips per line, ns p > 0.05, ** p < 0.005, one-way ANOVA). Values plotted as ± SEM. **(C)** Relative percentage of endogenous glutathione levels based on MCB fluorescence plotted out for each line tested (n = 3-6 fields of view per line; 2 coverslips per line, ns p > 0.05, * p < 0.05, ** p < 0.005, one-way ANOVA). Values plotted as ±SEM. **(D)** Quantification of the percentage of dead cells from control, and grouped familial SNCA PD lines (Control = 6.7% ± 1.2, SNCA PD = 13.0% ± 1.9) (n = 4-5 fields of view per line, 2 control lines, 2 A53T lines, and 1 SNCA x3 line, 1 neuronal induction, ** p = 0.0099, Welch’s t-test). Values plotted as ± SEM. **(E)** Representative images showing cells labelled with the nuclear marker Hoechst 33342 from control and SNCA PD lines. Arrows indicate pyknotic cells, defined by the small Hoechst 33342 positive dense cell area. Scale bar = 20μm. **(F) i)** Quantification of the percentage of pyknotic cells in control, A53T, and SNCA x3 hiPSC lines (Control = 14.9% ± 1.3, A53T = 18.3% ± 1.8, SNCA x3 = 21.8% ± 4.8), and **ii)** percentage of pyknotic cells in control, and grouped familial SNCA PD lines (Control = 14.9% ± 1.3, SNCA PD = 19.3% ± 1.8) (n = 4-5 fields of view per line, 2 control lines, 2 A53T lines, and 1 SNCA x3 line, 1 neuronal induction, ns: p = 0.059, Welch’s t-test). Values plotted as ± SEM.

